# Nuclear confinement from matrix stiffness drives epigenomic reprogramming of gingival fibroblasts

**DOI:** 10.64898/2026.05.27.728299

**Authors:** Hardik Makkar, Vinayak Vinayak, Kang I Ko, Rebecca G. Wells, Vivek B. Shenoy, Kyle H. Vining

## Abstract

Stromal fibroblasts reside in a three-dimensional (3D) extracellular matrix (ECM) that physically confines the cell and its nucleus. Nuclear mechanobiology studies have relied largely on two-dimensional substrates, on which cells spread freely without confinement. Therefore, it remains largely unknown how confinement in 3D stiff tissues regulates the fibroblast epigenome. Healthy human gingiva is characterized by a dense, stiff collagen ECM that undergoes softening in periodontal disease. A reduction in collagen in human gingiva correlated with elevated stromal H3K27 acetylation. In 3D hydrogels with stiffness matched to healthy and diseased gingiva, stiffness-dependent matrix confinement compacted the fibroblast nucleus, folded the nuclear envelope, and maintained low H3K27 acetylation through DNA methyltransferase activity, independently of actomyosin contractility. Stiff-matrix confinement increased DNA methylation at pericentromeric satellite repeats and decreased it at CTCF motifs. A stochastic chromatin model showed that broadening the range of chromatin contacts is sufficient to increase methylation in highly methylated regions and decrease it in less methylated regions, with changes concentrated at methylation-domain boundaries in both the model and the data. Stiff-matrix confinement also reduced CTCF occupancy uniformly across the genome, independent of CTCF expression and local DNA methylation. These changes led to hypomethylation of collagen gene promoter sequences, whereas hypermethylation of the MAP3K14 promoter attenuated non-canonical NF-κB inflammatory signaling. These findings establish stiffness-dependent 3D confinement as a physical regulator of the stromal epigenome and inflammatory states and motivate new mechanotherapies that restore tissue stiffness to resolve stromal cell states in chronic inflammatory diseases.

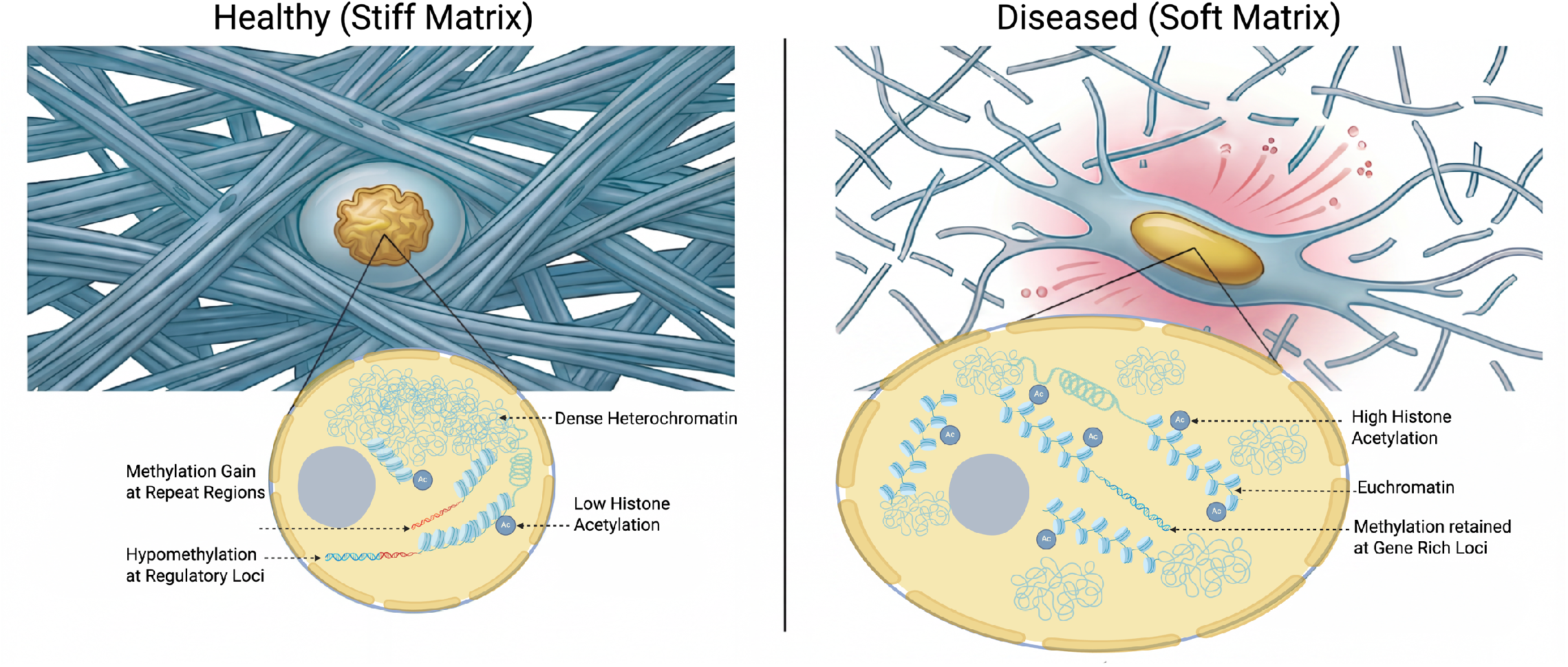
Graphical Abstract.

## Introduction

The extracellular matrix (ECM) surrounds cells in three dimensions (3D) and regulates their behavior through its mechanical properties and the physical confinement it imposes^1–3^. Stromal fibroblasts are subjected to this confinement in tissue homeostasis and disease^2–5^. These cells maintain connective tissue integrity across organs and regulate inflammatory responses through ECM synthesis and innate immune signaling. Studies of cell-ECM mechanotransduction have largely focused on cells cultured on two-dimensional (2D) elastic substrates of defined stiffness. Substrate stiffness directs stem cell lineage specification, scales nuclear lamin composition, and modulates DNA methylation and histone modifications^2^. In this geometry, however, cells spread laterally without mechanical resistance, and stiffness acts mainly by promoting cytoskeletal engagement^2,4,6,7^. In 3D, the surrounding ECM constrains cell volume and shape, alters nuclear morphology, and imposes compressive loads on the nuclear lamina^2^. The mechanisms by which mechanical cues regulate chromatin, therefore, differ between 2D and 3D environments. Defining how matrix stiffness regulates the stromal epigenome requires 3D matrices that match the stiffness of native tissue and impose the same constraints on cell and nuclear shape.

Gingival fibroblasts (GFs) are the principal stromal cells of the oral connective tissue. They produce the dense, architecturally organized fibrillar type I collagen networks in which they reside and coordinate innate immune responses to the resident oral microbiome. Gingival tissues heal exceptionally fast, with minimal scar formation, and tissue damage in periodontal disease rarely results in sepsis. In periodontal disease, proteases of host and microbial origin progressively degrade the ECM, lowering tissue stiffness and reducing the physical confinement that the healthy matrix imposes on resident fibroblasts^3,8–13^. Matrix degradation, tissue softening, and reduced fibroblast confinement are also shared features of musculoskeletal, ulcerative gastrointestinal, and other chronic inflammatory diseases. How stiff-matrix confinement, and its reduction in disease, regulates the chromatin and inflammatory states of stromal fibroblasts is unknown.

Isolating the effect of 3D confinement on the stromal epigenome requires a matrix in which stiffness-dependent confinement can be tuned independently of these variables. We used a hydrogel composed of interpenetrating networks of collagen and alginate^3,14,15^. Collagen concentration was kept constant across conditions, so cells encountered the same density of adhesion sites. Stiffness was tuned by the degree of crosslinking of alginate, which spans the storage modulus of healthy and diseased gingival tissue without altering polymer mesh architecture^3^. Alginate is the sole load-bearing network, and cellular remodeling does not alter the bulk stiffness of the hydrogel^3,14,15^. GFs in this system, therefore, experienced stiffness and 3D confinement as the primary mechanical variables.

Here, by pairing healthy and periodontitis-affected human gingival biopsies with donor-derived GFs in 3D hydrogels, we examined how stiffness-dependent confinement regulates fibroblast chromatin and inflammatory states. We combined nuclear imaging, transcriptomics, whole-genome bisulfite sequencing, CTCF CUT&RUN, and an in silico chromatin contact model, and tested whether these states persist after the mechanical environment changes. We show that stiff-matrix confinement compacts the nucleus, reduces H3K27 acetylation, redistributes DNA methylation, and reduces CTCF occupancy. Collagen genes are hypomethylated at their promoters and upregulated, whereas MAP3K14 promoter hypermethylation attenuates non-canonical NF-κB inflammatory signaling. These findings identify reduced confinement in soft matrices as a mechanism that increases stromal fibroblast responsiveness to inflammatory stimuli. This work supports the development of ECM-instructive therapies that restore healthy tissue mechanics in periodontitis and other matrix-degrading diseases.

## Results and Discussion

### 1. Collagen loss in periodontitis correlates with elevated H3K27Ac in the gingival stroma

We first asked whether collagen integrity in human gingival tissue is associated with the chromatin state of resident stromal cells. Healthy gingival connective tissue is characterized by dense, architecturally organized fibrillar collagen networks that confer its mechanical properties, which are lost in periodontitis-affected tissue due to protease-mediated collagen degradation and MMP-driven ECM remodeling^3,11^. In healthy gingival tissue, stromal fibroblasts display round, compact nuclei surrounded by dense fibrillar collagen detected by a second-harmonic generation (SHG) signal. In periodontitis-affected tissue, the SHG signal is lost alongside collagen degradation, and stromal nuclei appear elongated (**Fig. 1A**). SHG of fibrillar collagen architecture and immunofluorescence imaging of Acetyl-Histone H3 (Lys27) were performed on healthy and periodontitis-affected gingival biopsies to evaluate epigenetic markers of stromal cells in collagen matrix environments (**Fig. 1B**). H3K27Ac expression was significantly elevated in periodontitis-affected tissue with reduced SHG collagen signal compared to healthy samples (**Fig. 1C**). Spearman correlation across individual stromal cells established a significant negative relationship between collagen signal intensity and H3K27Ac level (rho = −0.732, **Fig. 1D**). Together, these data show that collagen loss in periodontitis is associated with elevated H3K27 acetylation in resident stromal cells.

**Figure 1.**
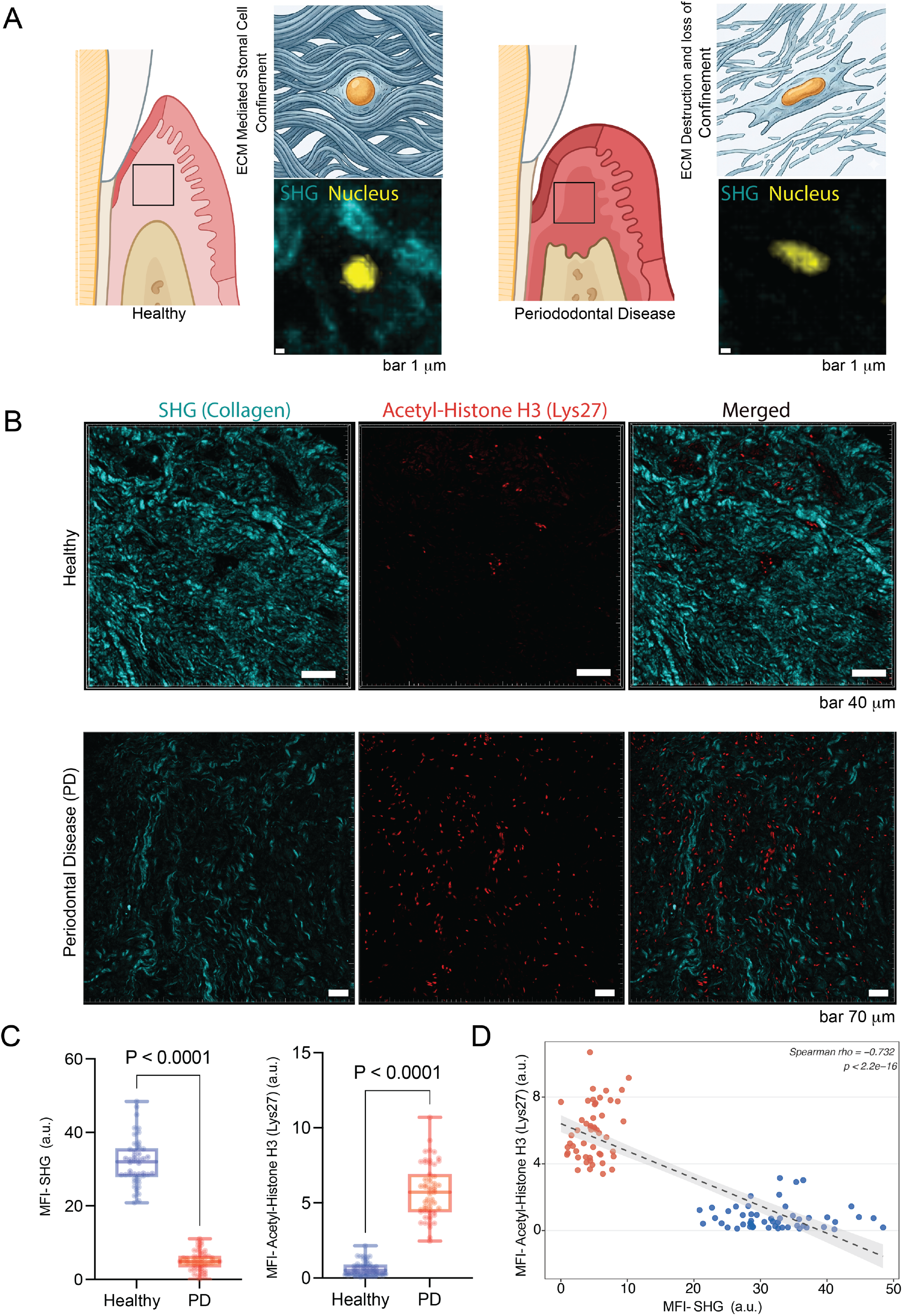
ECM collagen integrity correlates with stromal histone acetylation in human gingival tissues. (**A**) Schematic illustrating the relationship between ECM mechanical state and fibroblast nuclear morphology in healthy gingival tissue, where dense fibrillar collagen mediates stromal cell confinement, and in periodontal disease, where ECM degradation and loss of confinement accompany nuclear deformation. Representative SHG (cyan) and nuclear (yellow) images from healthy and periodontitis-affected tissue are shown. Scale bars, 1 µm. (**B**) Representative confocal immunofluorescence and second harmonic generation (SHG) images for Acetyl-Histone H3 (Lys27) (H3K27Ac, red) and fibrillar collagen (cyan) in healthy and periodontal disease (PD) human gingival tissue sections (n=3 biologically independent donors). Scale bars, 40 µm (healthy), 70 µm (PD). (**C**) Quantification of SHG mean fluorescence intensity (MFI) and H3K27Ac MFI in healthy and PD tissue. Data are shown as box plots. P-values determined by a two-tailed unpaired t-test. (**D**) Spearman correlation between SHG MFI and H3K27Ac MFI across individual stromal cells from healthy (blue) and PD (red) tissue sections. Dashed line indicates linear regression; shaded region, 95% confidence interval. Spearman rho = −0.732, p < 2.2×10⁻^16^.

### 2. Matrix stiffness governs nuclear geometry and nuclear envelope gene expression in gingival fibroblasts

Because collagen loss in human tissue tracked with elevated stromal H3K27Ac, we next asked whether matrix stiffness alone is sufficient to alter fibroblast nuclear geometry and chromatin state. We used a mechanically tunable hydrogel system to investigate how ECM stiffness-mediated confinement influences the epigenetic programming of stromal fibroblasts. ^3,16,17^ Gingival fibroblasts (GFs) isolated from healthy human donors were encapsulated within collagen-alginate interpenetrating network hydrogels. The storage modulus was tuned by ionic crosslink density to stiff (G’ 2.0 kPa) and soft (G’ 0.75 kPa), matching the rheological properties of human gingival tissue measured by shear rheology^3^ (**Fig. 2A, B**, and Supplementary **Fig. S1** and **Table 1**). Crosslink density was independent of ligand presentation and polymer mesh architecture^14,15,18^, ensuring that matrix stiffness and the confinement it imposes were the primary variables under investigation. Three-dimensional confocal reconstruction of F-actin and nuclear morphology (**Fig. 2C**) showed that GFs within soft (G’ 0.75 kPa) matrices exhibited morphologies ranging from moderate elongation (aspect ratio 1.70) to pronounced bipolar extension (aspect ratio 3.21). Encapsulation within the stiff (G’ 2.0 kPa) matrix produced a confined, near-isotropic nuclear geometry (aspect ratio 1.15). Quantification across biological replicates confirmed that nuclear aspect ratios were significantly lower and less dispersed in the stiff (confined) matrices (**Fig. 2D**), with a broader interquartile range in the soft condition, indicating that matrix compliance is associated with a heterogeneous landscape of nucleo-cytoskeletal configurations rather than a single deformation state^19^. Nuclear volume was influenced primarily by matrix stiffness and actomyosin contractility (**Fig. 2E**). Inhibition of non-muscle myosin II (Blebbistatin) or ROCK (Y-27632) in soft-matrix prevented cell spreading cells significantly reduced nuclear volume, whereas there was no effect on cells in stiff matrix. (Supplementary **Fig. S1 D,E)**. This suggests that confined nuclear geometry in a stiff matrix is independent of contractile status.

**Figure 2.**
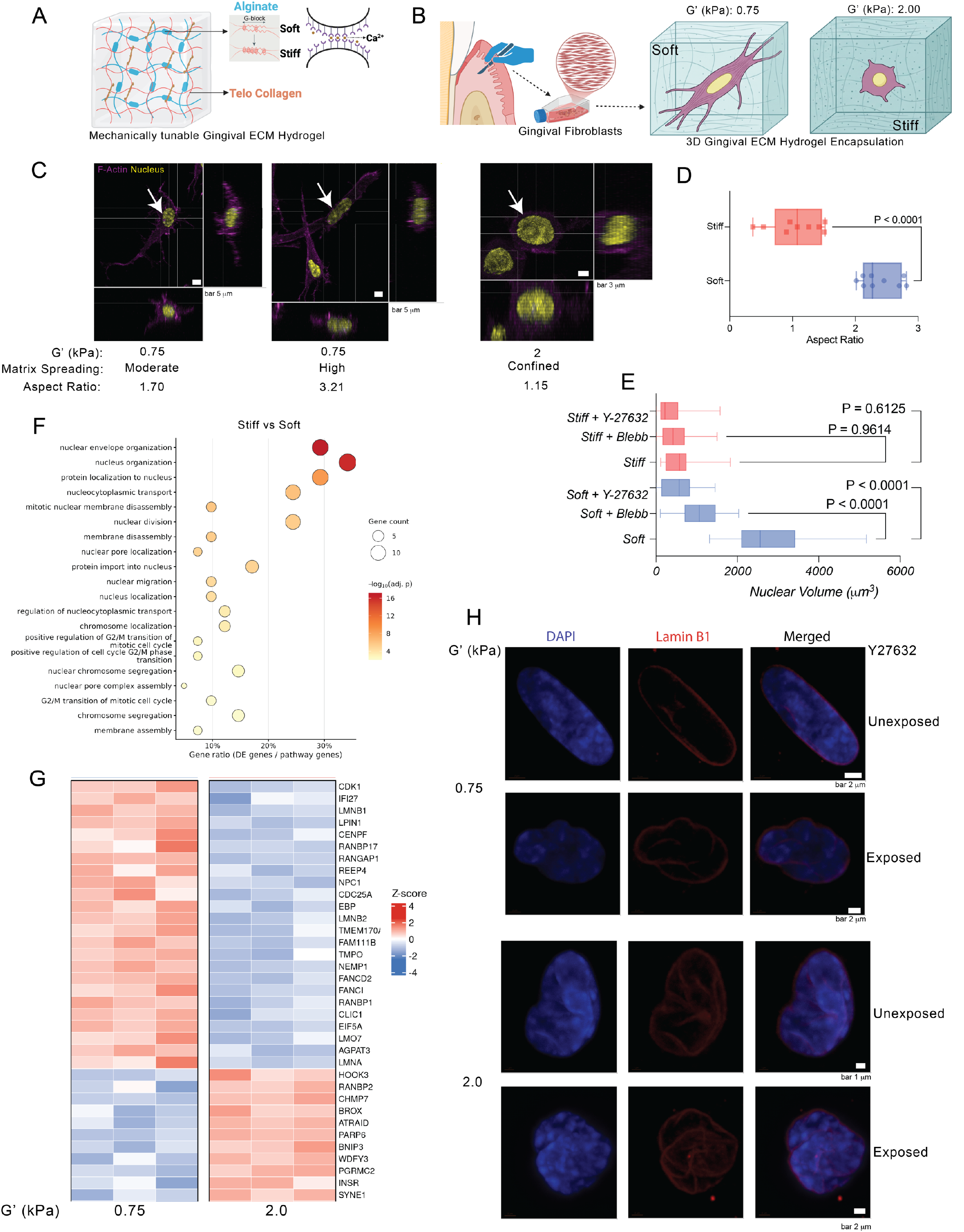
Three-dimensional matrix confinement enforces isotropic nuclear geometry, drives divergent lamin expression, and produces mechanistically distinct nuclear envelope architectures. (A) Schematic of the collagen-alginate interpenetrating network (IPN) hydrogel system comprising of ionic crosslinking of the alginate network and type I telo-collagen. (B) Schematic illustrating GF encapsulation in soft (G’ 0.75 kPa) and stiff (G’ 2.0 kPa) 3D gingival ECM hydrogels. (C) Representative 3D confocal immunofluorescence images of F-actin (magenta) and nucleus (yellow) in GFs encapsulated in soft and stiff hydrogels, showing three characteristic morphologies. Aspect ratios are indicated. Scale bars, 5 µm (soft), 3 µm (stiff). (D) Quantification of nuclear aspect ratio in soft and stiff hydrogels. Data shown as box plots. P-value determined by two-tailed unpaired t-test. (E) Nuclear volume quantification across stiffness conditions and following treatment with the myosin II inhibitor Blebbistatin (Blebb) or the ROCK inhibitor Y-27632. Data shown as box plots. P-values determined by two-way ANOVA with Šídák’s multiple comparisons test. (F) Bubble plot showing Gene Ontology biological process enrichment for differentially expressed genes in stiff versus soft hydrogels. Bubble size indicates gene count; colour scale indicates −log_10_(adjusted p-value). (G) Heatmap showing z-scored expression of nuclear envelope and transport-related genes in GFs in soft (G’ 0.75 kPa) and stiff (G’ 2.0 kPa) hydrogels from bulk RNA sequencing (n = 3). (H) Representative confocal immunofluorescence images of DAPI (blue) and Lamin B1 (red) in GFs in soft (G’ 0.75 kPa) and stiff (G’ 2.0 kPa) matrices, unexposed or exposed to Y-27632. Scale bars as indicated. P values <0.05 were considered statistically significant and are indicated on the graphs. Panels A and B created with Biorendor.com

Transcriptomic profiling by bulk RNA sequencing revealed a stiffness-dependent regulation of nuclear lamina composition (**Fig. 2G**). B-type lamins form the primary structural scaffold of the inner nuclear lamina in proliferating cells. *LMNB1* and *LMNB2* were elevated in GFs in the soft matrix. They were co-expressed with the mitotic regulatory genes *CDK1, CDC25A, and CENPF*, the inner nuclear membrane Lamin B-binding protein TMPO, encoding the lamina-associated polypeptide LAP2, and the nuclear envelope remodeling factor *NEMP1*. This suggests a mechanically permissive cell state, in which a lamina enriched in compliant B-type lamins accommodates cytoskeletal force-driven nuclear deformation^20–22^. Lamin A/C levels scale with ECM stiffness through a tension-dependent post-translational stabilization mechanism.^22^ *LMNA* was lower in GFs encapsulated in a stiff matrix. Nesprin-1 is the outer nuclear membrane LINC complex component that bridges the actomyosin cytoskeleton to the nuclear lamina. *SYNE1* was elevated in GFs encapsulated in stiff matrix, pointing toward reinforcement of the nuclear-cytoskeletal connections required for force transmission to the nuclear interior^23,24^. The upregulation of nucleocytoplasmic transport regulators *RAN, RANBP2, KPNB1, CSE1L, and HOOK3* in the stiff condition is consistent with increased nuclear import demands ^20^. Gene ontology analysis showed nuclear envelope organization was the most significantly enriched biological process in the stiff-versus-soft comparison, followed by nucleus organization, nucleocytoplasmic transport, protein localization to nucleus, and nuclear pore complex assembly (**Fig. 2F**). The transcriptional reinforcement of nuclear envelope and transport machinery under confinement also has consequences for the nuclear entry of mechanosensitive transcription factors. YAP, a central mechanotransducer whose activity depends on nuclear import^25^, showed partial nuclear translocation in stiff-matrix GFs relative to soft (**Fig. S2A, B**). YAP/Hippo pathway gene loci carried predominantly hypomethylated DMRs in the stiff condition (**Fig. S2C,D**), indicating that these mechanosensory genes remain accessible despite confinement.

### Actomyosin contractility and matrix stiffness produce distinct nuclear envelope folding architectures

Having established that stiff matrix alters nuclear geometry and lamin gene expression, we next asked whether stiffness also changes nuclear envelope topology, and whether this change depends on actomyosin contractility. Lamin B1 immunostaining, used as a structural reporter of nuclear envelope topology, revealed that GFs in soft matrix displayed smooth, geometrically regular nuclear envelopes sustained by cytoskeletal pre-stress (**Fig. 2H**). Inhibition of ROCK-dependent actomyosin contractility in soft-matrix showed pronounced nuclear envelope wrinkling and folding in GFs, consistent with latent buckling instability of this pre-stressed lamina^21^. ROCK inhibition in GFs in stiff matrices left fold morphology unchanged, indicating that confinement-driven folding is independent of actomyosin contractility, likely driven by compressive loads from the crosslinked polymer network ECM. Fold morphologies were heterogeneous among individual confined nuclei, with spatial distribution, angular orientation, and amplitude varying substantially under the same conditions, reflecting the stochastic distribution of local compressive stresses within the gingival ECM hydrogel.

Structure tensor coherence analysis of Lamin B1 terrain maps computed the coherence index C = (λ₂ − λ₁)/(λ₂ + λ₁) from the local intensity gradient tensor, where a score of 0 indicates an isotropic gradient with no preferred fold orientation, and 1 indicates a perfectly linear fold axis (**Supplementary Fig. S3**). Soft-matrix nuclei exhibited high coherence (0.7045), with bilateral polar lobe profiles aligned with the nuclear long axis. Stiff-matrix nuclei showed lower coherence (0.4569), with polar profiles fragmenting into multiple low-amplitude spikes distributed across all angular positions. We propose that the distinct folding architecture reflects differential engagement of the chromatin– and lamin-governed mechanical regimes of the nucleus ^21,26^, driven by the mechanically distinct environments of soft and stiff matrices. Together, these data identify two mechanically distinct routes to nuclear envelope folding. In a soft matrix, ROCK-mediated actomyosin tension holds the lamina under pre-stress, and releasing this tension induces folding. In a stiff matrix, folding is unaffected by ROCK inhibition. This points to stiffness-induced matrix confinement, rather than cytoskeletal tension, as the source of lamina compression.

### 3. Matrix compliance drives H3K27Ac accumulation through actomyosin and DNA methyltransferase-dependent nuclear deformation in GFs

Having established that nuclear volume in a soft matrix depends on actomyosin contractility, whereas compact, folded nuclei in a stiff matrix do not respond to contractility inhibition, we next asked whether these nuclear states regulate active enhancer marks. We examined H3K27Ac, which marks active enhancers and promoters, across stiffness conditions with or without inhibition of actomyosin contractility. Actomyosin contractility regulates histone deacetylase nucleocytoplasmic shuttling and histone acetyltransferase nuclear import in fibroblasts^27,28^. GFsin soft matrices exhibited high H3K27Ac MFI that was significantly attenuated by both Blebbistatin and Y-27632, converging toward the low-acetylation state of confined cells in stiff matrices (**Fig. 3A, B**). In stiff matrices, H3K27Ac was constitutively low and remained unchanged upon actomyosin contractility inhibition. Nuclear volume and H3K27Ac MFI co-varied across all conditions (**Fig. 2E, 3B**). Actomyosin inhibition reduced nuclear volume and H3K27Ac expression in soft matrix. In the stiff matrix, neither nuclear volume nor H3K27Ac expression responded to inhibitor treatment. The compacted nuclear geometry under 3D confinement likely limits acetyltransferase access via steric hindrance rather than enzymatic regulation ^28^, because chromatin condensation states dictate both the mechanical response and the physical accessibility of the genome^26^. These data highlight the impact of nuclear deformation on the stromal epigenome in soft ECM conditions (**Fig. 3C**). To determine whether this mechano-epigenomic coupling is a general property of stromal fibroblasts rather than a tissue-specific response, we examined BJ dermal fibroblasts under identical stiffness conditions. Stiffness-dependent regulation of H3K27Ac and nuclear Lamin B1 folding was preserved in BJ cells, with soft-matrix fibroblasts maintaining high H3K27Ac that was attenuated by Y-27632 and stiff-matrix fibroblasts exhibiting constitutively low H3K27Ac and compacted, folded nuclei insensitive to contractility inhibition (Supplementary **Fig. S4).**

**Figure 3.**
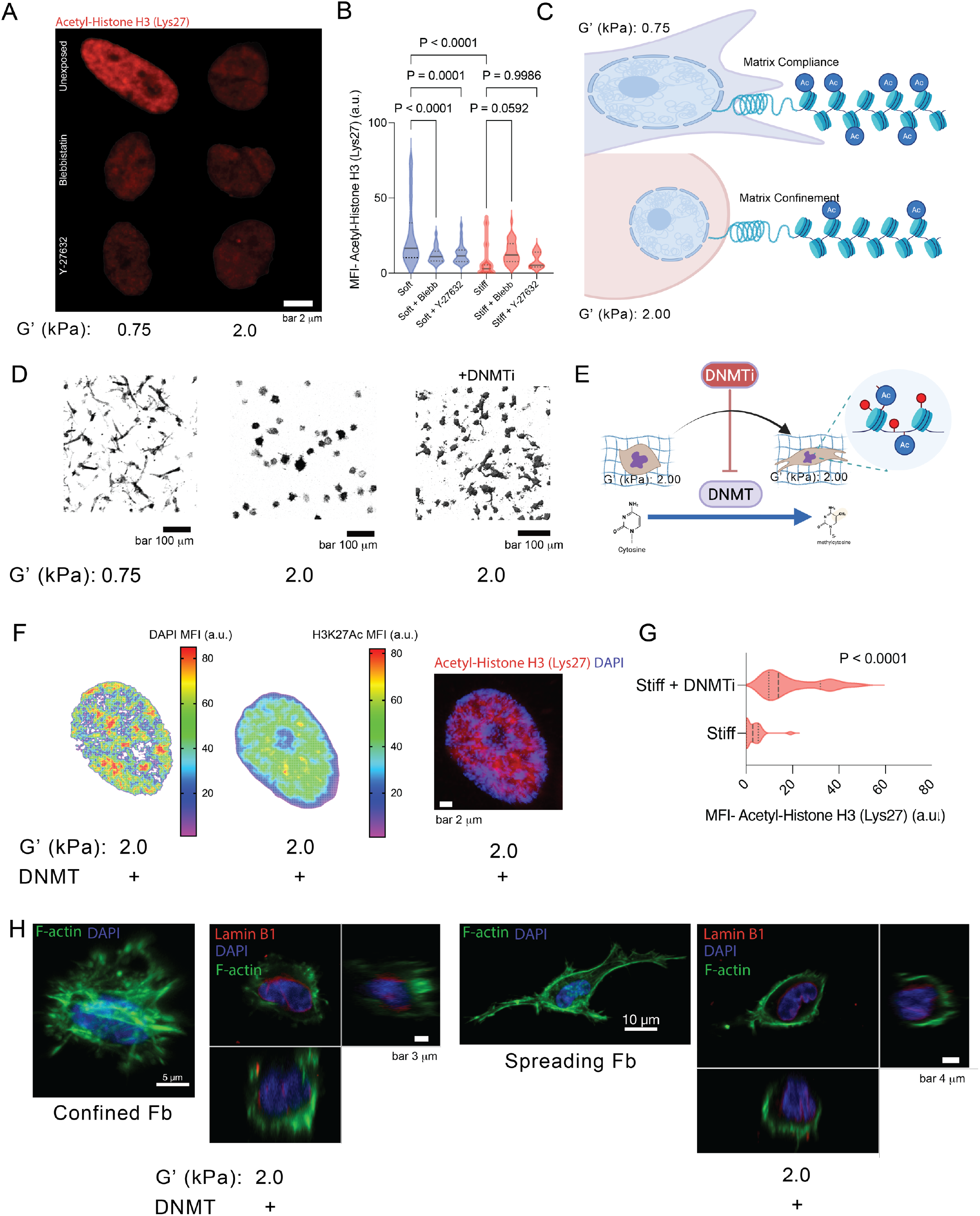
ECM stiffness suppresses H3K27Ac through actomyosin-dependent nuclear geometry and DNMT1 activity, with persistent nuclear envelope folding following epigenetic reversal. (A) Representative confocal immunofluorescence images of Acetyl-Histone H3 (Lys27) (H3K27Ac, red) in GFs encapsulated in soft (G’ 0.75 kPa) and stiff (G’ 2.0 kPa) hydrogels, untreated or treated with Blebbistatin or Y-27632. Scale bar, 2 µm. (B) Quantification of H3K27Ac MFI across all conditions. P-values determined by two-way ANOVA with Šídák’s multiple comparisons test. (C) Schematic summarizing the relationship between matrix compliance, actomyosin contractility, and histone acetylation in soft and stiff matrices. (D) Representative z-projection confocal images of phalloidin-stained GFs in soft (G’ 0.75 kPa), stiff (G’ 2.0 kPa), and stiff+DNMTi matrices. Scale bars, 100 µm. (E) Schematic illustrating DNMT inhibitor (DNMTi) treatment in stiff matrices and its effect on cytosine methylation. (F) DAPI MFI heatmap and H3K27Ac MFI heatmap of representative nuclei from GFs in stiff matrices treated with DNMTi, with merged immunofluorescence image showing H3K27Ac (red) and DAPI (blue). Scale bar, 2 µm. (G) Quantification of H3K27Ac MFI in stiff and stiff+DNMTi conditions. Data shown as violin plots. P-value determined by two-tailed unpaired t-test. (H) Representative confocal immunofluorescence images of F-actin (green), DAPI (blue), and Lamin B1 (red) in confined fibroblasts (stiff, G’ 2.0 kPa, DNMTi treated) and DNMTi-induced spreading fibroblasts (stiff, G’ 2.0 kPa, DNMTi treated). Lamin B1 wrinkling persists in the spreading fibroblast despite morphological reversal. Scale bars as indicated. Panel C and E created with Biorendor.com

DNMTi treatment of GFs in the stiff matrix produced a marked reorganization of the nuclear chromatin (**Fig. 3D, E**). H3K27Ac heatmaps shifted from the low-signal, spatially uniform distribution of the confined state to a broad, high-intensity acetylation pattern, with H3K27ac mean fluorescence intensity (MFI) increasing substantially compared to the untreated stiff condition (**Fig. 3F, G**). DAPI intensity maps in soft matrix showed a hotspot-dominated heterochromatin pattern. These results were consistent with our previous findings that DNMTi restored cell spreading and nuclear volume expansion in the stiff matrix^3^. The loss of DNA methylation,^29^ recovery of H3K27ac, and structural reorganization of chromatin under DNMTi treatment suggest the hypothesis that CpG methylation at regulatory loci restricts acetyltransferase recruitment by limiting physical chromatin accessibility.^30^ GFs in stiff matrices showed a highly compacted, folded nuclear envelope morphology (**Fig. 3H**). Following DNMTi treatment, although the fibroblasts began to spread, morphologically resembling cells in soft matrices, nuclear wrinkling persisted rather than resolving into the smooth envelope architecture characteristic of untreated GFs in soft matrices. This indicates that the nuclear envelope topology established under confinement persists longer than the initial epigenetic and morphological reversal produced by DNMTi. Previous studies show that 3D confinement and HDAC-driven chromatin condensation can permanently alter nuclear mechanics and lamin distribution in fibroblasts, leaving structural changes that persist even after the cells are removed from the confining environment or the matrix is softened^5,6^. Whether this retained Lamin B1 fold topology reflects a confinement-imprinted mechanical memory remains undetermined. Validating this hypothesis will require time-resolved imaging of lamin dynamics and lamina-associated domain (LAD) repositioning during the DNMTi-induced morphological changes. Together, these data identify two mechanically distinct modes of H3K27Ac regulation in GFs. In compliant matrices, H3K27Ac scales with actomyosin-dependent nuclear expansion, and both are attenuated by inhibition of myosin II or ROCK. Under stiff matrix confinement, compacted nuclei maintain a low-acetylation state that is refractory to contractility inhibition and relieved by DNMT inhibition. Lamin B1 folding persists despite the DNMTi-induced increase in H3K27Ac and cell spreading, which indicates that nuclear envelope topology and chromatin acetylation are decoupled during epigenetic reversal.

### 4. Matrix stiffness drives redistribution of H3k9me3 and opposing methylation states at repetitive and regulatory genomic loci in GFs

We next asked whether matrix stiffness reorganizes constitutive heterochromatin and the genome-wide DNA methylome. To test this, we compared untreated GFs in soft and stiff matrices by H3K9me3 imaging and whole-genome bisulfite sequencing (WGBS). Constitutive heterochromatin organization was assessed by plotting DAPI and H3K9me3 fluorescence intensity heatmaps to determine whether matrix stiffness alters the spatial distribution of the repressive chromatin compartment in gingival fibroblasts (**Fig. 4 A,B**). Soft-matrix nuclei displayed discrete DAPI-dense heterochromatin foci with punctate, co-localized H3K9me3 signal, whereas these signals redistributed into a diffuse, spatially homogeneous nuclear pattern of GFs in stiff-matrix. Normalized H3K9me3 intensity profiles from biological replicates are shown in Supplementary **Fig. S5**. Soft-matrix nuclei generated high-amplitude, multi-peak profiles reflecting spatially clustered heterochromatin (mean CV ∼58%), while stiff-matrix nuclei produced broad plateau-like profiles (mean CV ∼25%). The redistribution of H3K9me3 expression in GFs from focal clusters toward a spatially diffuse nuclear distribution under stiff conditions is consistent with nuclear deformation during cell migration through confined spaces^31^. Our results show that stiff ECM induces a stable structural adaptation in GFs to mechanical confinement. This is also consistent with past work showing the loss of H3K9me3 and nuclear softening under transient mechanical stretch, which protects the genome from mechanical damage^32^.

**Figure 4.**
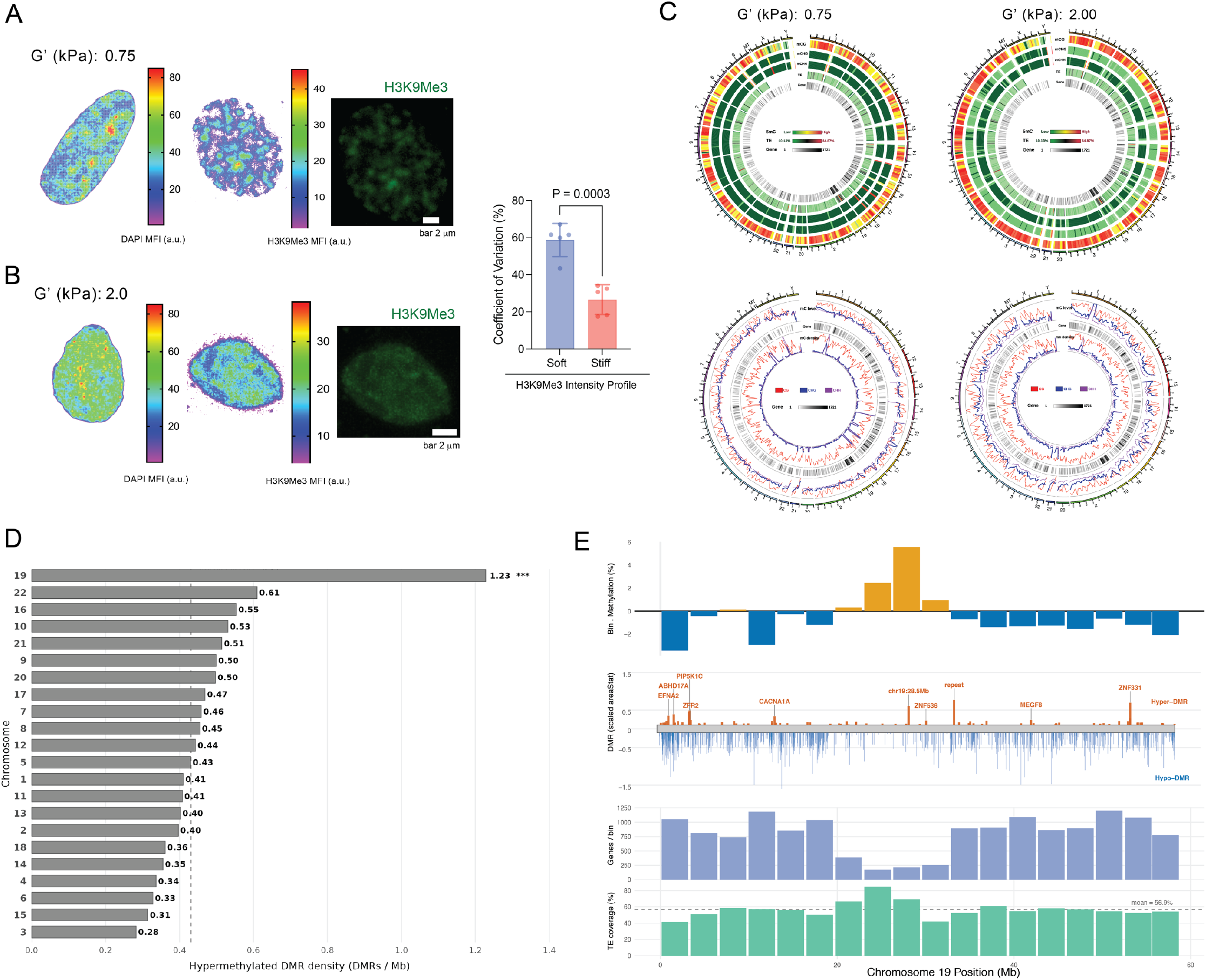
Matrix confinement drives heterochromatin redistribution, and a bifurcated methylation program targeting pericentromeric repeats and preserving regulatory chromatin. (A,B) Representative DAPI MFI heatmaps, H3K9me3 MFI heatmaps, and H3K9me3 immunofluorescence images of GF nuclei in soft (G’ 0.75 kPa) and stiff (G’ 2.0 kPa) matrices. Scale bars, 2 µm. Bar graph shows coefficient of variation (CV) of H3K9me3 intensity profiles across conditions (n = 5 biological replicates). P-value determined by two-tailed unpaired t-test. (C) Genome-scale circos plots showing CpG methylation context (mCG) density across all chromosomes in GFs from soft (G’ 0.75 kPa) and stiff (G’ 2.0 kPa) matrices. (D) Bar chart showing hypermethylated DMR density (DMRs per Mb) across all autosomes. Chromosome 19 carries the highest density (1.23 DMRs/Mb) One sided Poisson Exact Test with Benjamini-Hochberg correction; p_adj (BH) 5.3×10^−15^ (E) High-resolution genomic landscape of chromosome 19, showing from top to bottom: bin-level methylation percentage (orange, hypermethylated bins; blue, hypomethylated bins); DMR track with scaled area statistics (orange, Hyper-DMRs; blue, Hypo-DMRs) with selected gene annotations; genes per genomic bin; and transposable element (TE) coverage percentage with chromosomal mean indicated

Next, we performed whole-genome bisulfite sequencing to determine a stiffness-associated reorganization of the GFs methylome (**Fig S6A, B,C**). Genome-scale circos plots visualized CpG cytosine methylation states (mCG) (**Fig. 4C**). Genome-scale circos plots showed higher methylated-CpG density in the stiff condition across large genomic bins (**Fig. 4C**). This density measure reflects regional methylated-CpG concentration rather than a direct locus-level comparison between conditions. Elevated DNMT1 nuclear expression in the stiff condition has been documented previously^3^ and no differences were observed in DNMT3 expression between soft and stiff matrices at transcript and protein levels (**Fig S7A,B,C**). Together with prior reports of stiffness-associated DNA methylation changes in stromal cells^28,33,34^, these results are consistent with regional methylation gain at these loci.

Genome-wide analysis of the Δβ distribution across all autosomes confirmed that heterochromatin-associated loci predominantly gained methylation in the stiff condition, while euchromatin-associated loci showed predominantly neutral or negative methylation changes across every chromosome (**Fig. S8A**). The genomic feature distribution of DMRs resolved this bifurcation quantitatively. Hyper-DMRs were heavily skewed toward repeats (61%), with introns (18%), CpG islands/shores (9%), and promoters/TSS (7%) accounting for the remainder. Hypo-DMRs showed a relatively greater representation at promoters/TSS (12%), CpG islands/shores (13%), and introns (24%), with repeats accounting for 46% (**Fig. S8B**). This shows that methylation gain preferentially targets repeat-dense regions, while methylation loss is concentrated at regulatory sequences, including promoters and CpG islands. Further, chromosomal mapping of hypermethylated DMR density identified chromosome 19 as the primary condensation locus, carrying the highest hypermethylated DMR density of any autosome (**Fig. 4D,E and Fig. S9A,B**). Chromosome 19 analysis revealed that hypermethylated bins overlapped with transposable element-dense regions and pericentromeric repeats (mean TE coverage 56.9%), while hypomethylated bins were broad, gene-dense intervals containing regulators including *ZNF536*, *ZNF331*, *MEGF8, CACNA1A, and EFNA2*.

Among repeat classes, satellite repeats, the primary structural constituents of pericentromeric and telomeric constitutive heterochromatin, showed the greatest and most consistent methylation gain in the stiff condition (**Fig. 5A**). Satellite repeat methylation gain connects the methylation changes to the H3K9me3 redistribution (Fig. 4A). Satellite repeat transcription is associated with chromosomal instability, and its preferentially enhanced methylation under confinement is interpreted as a genome-stability response to sustained matrix stiffness-mediated confinement^32,35,36^. Differential methylation analysis at single-locus resolution revealed a dominant CG hypomethylated DMR population distributed across every chromosomal arm (**Fig. 5B**) in the stiff condition. Hypomethylated DMRs in the stiff condition were analyzed for DNA binding motif enrichment to determine whether the regulatory sequences that lose methylation under confinement are enriched for factors with defined roles in chromatin architecture and gene regulation (**Fig. 5C**). CTCF motifs were among the most significantly enriched. CTCF binding requires unmethylated CpG at its recognition sequence, and its enrichment at hypomethylated loci in the stiff condition indicates that CTCF binding sites are selectively demethylated under confinement. CTCF is a chromatin architectural protein that anchors chromatin loops and maintains topological domain boundaries across the mammalian genome^37–39^. Whether these demethylated sites correspond to active CTCF-bound loop anchors in confined fibroblasts remains to be established by binding and chromatin conformation assays. BORIS occupies overlapping recognition sequences with distinct outcomes, and its co-enrichment suggests the relative CTCF and BORIS occupancy at shared sites may differ between conditions. HINFP and Zfp57 enrichment connect these hypomethylated loci to active chromatin assembly and imprinting-associated regulatory sequences, respectively^40^.

**Figure 5.**
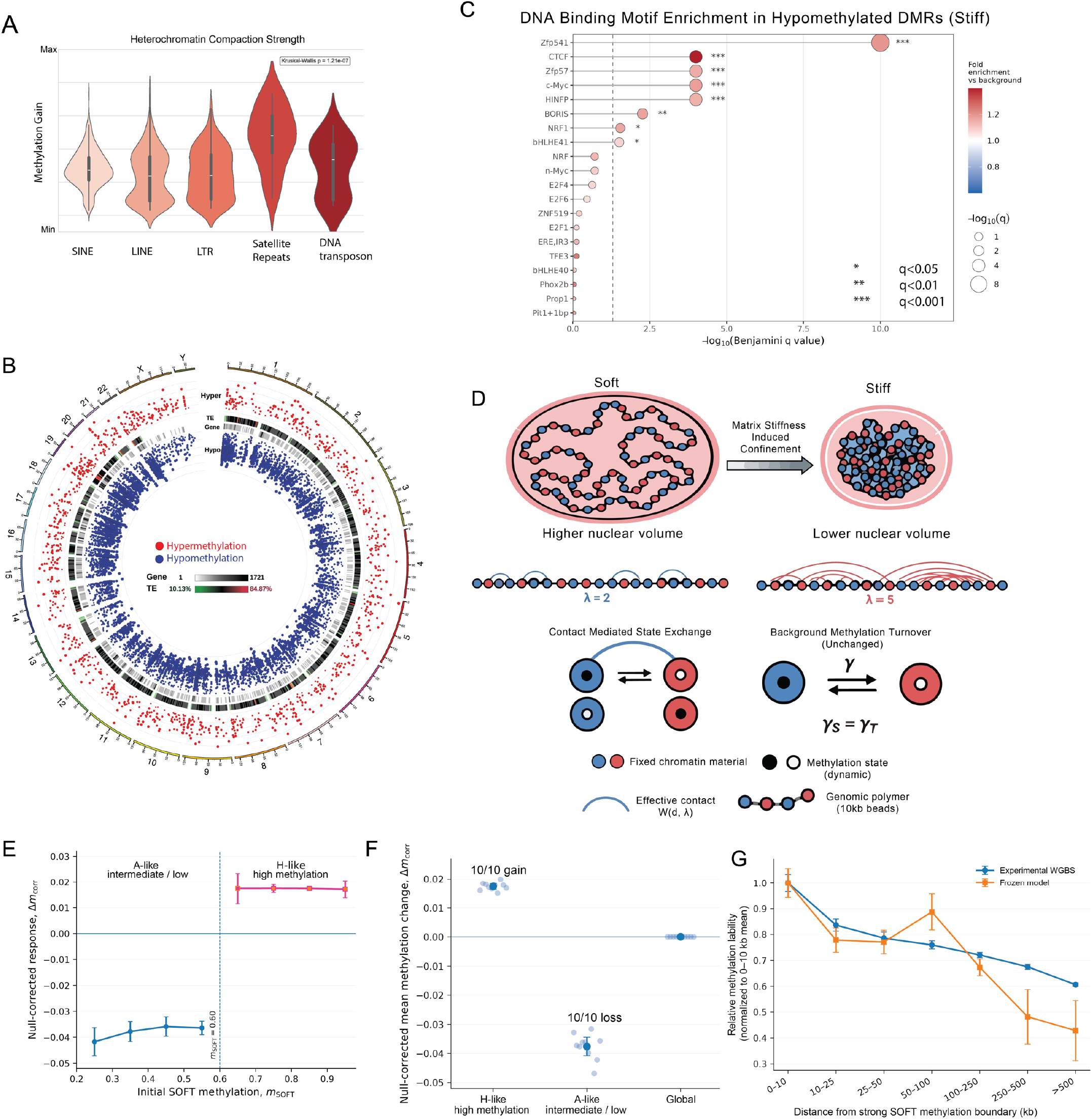
Matrix confinement drives a bifurcated methylation program across repressed and regulatory genomic loci. (A) Violin plots showing methylation gain across transposable element classes in the stiff condition relative to soft. Kruskal-Wallis p = 1.21×10⁻⁷. (B) Circos plot showing the genome-wide distribution of hypermethylated (red) and hypomethylated (blue) differentially methylated regions (DMRs) in the stiff condition. (C) Bubble plot showing transcription factor binding site motif enrichment within hypomethylated DMRs in the stiff condition. Bubble size indicates −log₁₀(q-value); colour scale indicates fold enrichment versus background. Significance thresholds: * q < 0.05, ** q < 0.01, *** q < 0.001. (**D**) Coarse-grained computational model linking nuclear confinement to methylation redistribution. The experimentally motivated soft matrix methylation landscape is represented at 10-kb resolution. Soft matrix chromatin is modeled with shorter-range effective genomic interactions (λ= 2), whereas stiff matrix confinement broadens and increases the density of effective chromatin contacts (λ= 5); the corresponding integrated one-sided contact weights are 1.00 and 2.87. The genomic sequence and initial methylation architecture are unchanged between conditions, together with the cooperative interaction strength, material fields, and background methylation-turnover rate. Only the confinement-dependent effective contact landscape is changed. Full model formulation and parameters are provided in the Supplementary Methods and Supplementary Table T4. (**E**) Relationship between the initial soft matrix methylation state and the null-corrected confinement-induced methylation response across modeled 10-kb bins. For each of 10 independently generated empirical-like soft matrix landscapes, bin-wise confinement responses were averaged within fixed initial-methylation intervals. The vertical dashed line at mSOFT = 0.60 is the pre-specified division between A-like intermediate/low methylation material and H-like high-methylation material. Confinement drives predominantly negative changes below this threshold and positive changes above it. **(F)** Mean methylation change in H-like high-methylation material, A-like intermediate/low-methylation material, and genome-wide across 10 independently generated empirical-like soft matrix landscapes. Individual points represent independent landscapes; summary markers and error bars indicate the across-landscape mean and 95% confidence interval. H-like regions reproducibly gain methylation (Δm= +0.0175, 95% CI +0.0165 to +0.0184), whereas A-like regions reproducibly lose methylation (Δm=−0.0377, 95% CI −0.0410 to−0.0345). The genome-wide mean methylation shift remains approximately zero, demonstrating redistribution rather than a uniform global change. The H/A classes are defined from initial SOFT methylation state and are not genomic-feature annotations. (**G**) Boundary-proximal methylation lability in experimental WGBS data and the frozen model. Strong boundaries were defined from the SOFT methylation landscape using adjacent methylation-state domains with both flanks >= 20 kb and a mean methylation difference >= 0.15. For WGBS, mean absolute STIFF-SOFT methylation change was calculated as a function of genomic distance from the nearest strong SOFT boundary. For the model, the same boundary-distance analysis was applied to the control-corrected confinement response. Model points show the mean across 10 independently generated SOFT landscapes, with uncertainty summarized across landscapes. Model boundaries are methylation-state boundaries and are not assigned from CTCF occupancy or experimentally measured topological domains.

### 5. Increasing the effective contact range generated a state-dependent redistribution of methylation across the pre-existing SOFT methylation landscape

To test how confinement-driven changes in chromatin interactions generate the opposing methylation responses observed experimentally, we developed a coarse-grained stochastic model of chromatin methylation^38,41,42^ (**Fig. 5D**). The genome was represented at 10-kb resolution using the heterogeneous methylation architecture measured in the SOFT condition. Highly methylated domains (m≥ 0.60) were represented as H-like, high-methylation/methylation-permissive chromatin material, whereas intermediate– and low-methylation domains (m < 0.60) were represented as A-like chromatin material. These material classes were defined from the soft methylation state. The model, therefore, does not attempt to generate the baseline methylome de novo but instead asks how an established heterogeneous methylation landscape responds to altered confinement. Matrix confinement was represented by increasing the effective genomic range and density of chromatin contacts while holding the initial methylation architecture, cooperative interaction strength, material fields, and background methylation turnover fixed. The effective interaction length increased from λ = 2 in soft to λ = 5 in stiff conditions, corresponding to an approximately 2.87-fold increase in integrated contact weight. Full model definition, parameterization, stochastic update rules, and sensitivity analyses are provided in the Supplementary Methods and Supplementary **Table T4**; preservation of the SOFT reference state over the finite 1-MCS readout is shown in Supplementary **Fig. S10**.

When modeled 10-kb bins were grouped by their initial soft methylation state, confinement produced negative methylation changes across intermediate/low-methylation regions and positive changes across highly methylated regions (**Fig. 5E**). Consistent with this state-dependent response, across 10 independently-generated empirical soft landscapes, H-like high-methylation regions consistently gained additional methylation under the stiff contact regime (Δm= +0.0175, 95% CI +0.0165 to +0.0184), whereas A-like intermediate/low-methylation regions consistently lost methylation (Δm=−0.0377, 95% CI−0.0410 to−0.0345; **Fig. 5F**). The expected direction was observed in all 10 independent landscapes. In contrast, the genome-wide mean methylation shift was only +7.6 × 10−5 (95% CI +6.4 × 10−5 to +8.8 × 10−5), indicating that confinement primarily produces regional redistribution rather than a uniform shift in global methylation (**Fig. 5F**) (**Fig. S10-13**). Because H/A identity was not assigned from genomic annotations, these responses should not be interpreted as direct predictions of repeat-, heterochromatin-, euchromatin-, or regulatory-locus methylation. Instead, the model demonstrates that broadening the effective contact landscape is sufficient to generate opposing methylation responses across a pre-existing heterogeneous methylation architecture while leaving the genome-wide mean nearly unchanged.

### 6. Methylation changes are concentrated near pre-existing methylation-domain bound areas

Methylation changes were concentrated near pre-existing SOFT methylation-state boundaries in both the experimental WGBS data and the model **(Fig. 5G**). Experimentally, mean absolute STIFF-SOFT methylation change was greatest near strong SOFT boundaries and decreased with genomic distance toward domain interiors. The frozen model reproduced this boundary-proximal enrichment and spatial decay. Relative to an independent SOFT-contact null branch, the model near-boundary methylation-lability excess was +0.00852 (95% CI +0.00734 to +0.00970). These boundaries represent transitions between methylation states in the coarse-grained model and are not assigned from CTCF occupancy or experimentally measured topological-domain boundaries. Their preferential lability is therefore not a direct prediction of CTCF binding, although it is consistent with the experimental enrichment of CTCF motifs among hypomethylated DMRs. The directional model response was reproduced across 20 independent master seeds and remained robust to variation in cross-material coupling, cooperative interaction strength, and background turnover (**Supplementary Figs. S11, S12, S14, and S15**). Together, the simulations support a physical mechanism in which nuclear confinement broadens the effective chromatin-contact landscape and is sufficient to redistribute methylation across a pre-existing heterogeneous methylation architecture.

### 7. Matrix confinement uniformly reduces CTCF occupancy in GFs

CTCF motifs were enriched within hypo-DMRs in the stiff condition (**Fig. 5C**, **Fig. 6A**), but whether these sites are CTCF-bound in GFs, and whether confinement alters CTCF binding, was not known. CTCF is the principal architectural protein that anchors chromatin loops with cohesin and marks the boundaries of topologically associating domains (TADs)^37^. CTCF binding sites also mark the borders of lamina-associated domains (LADs), the chromatin associated with the nuclear lamina^43^, which folds under confinement (**Fig. 2H**). We examined CTCF occupancy across the genome, at TAD boundaries, and at LAD borders in GFs in soft (G′ 0.75 kPa) and stiff (G′ 2.0 kPa) matrices by CUT&RUN.

**Figure 6.**
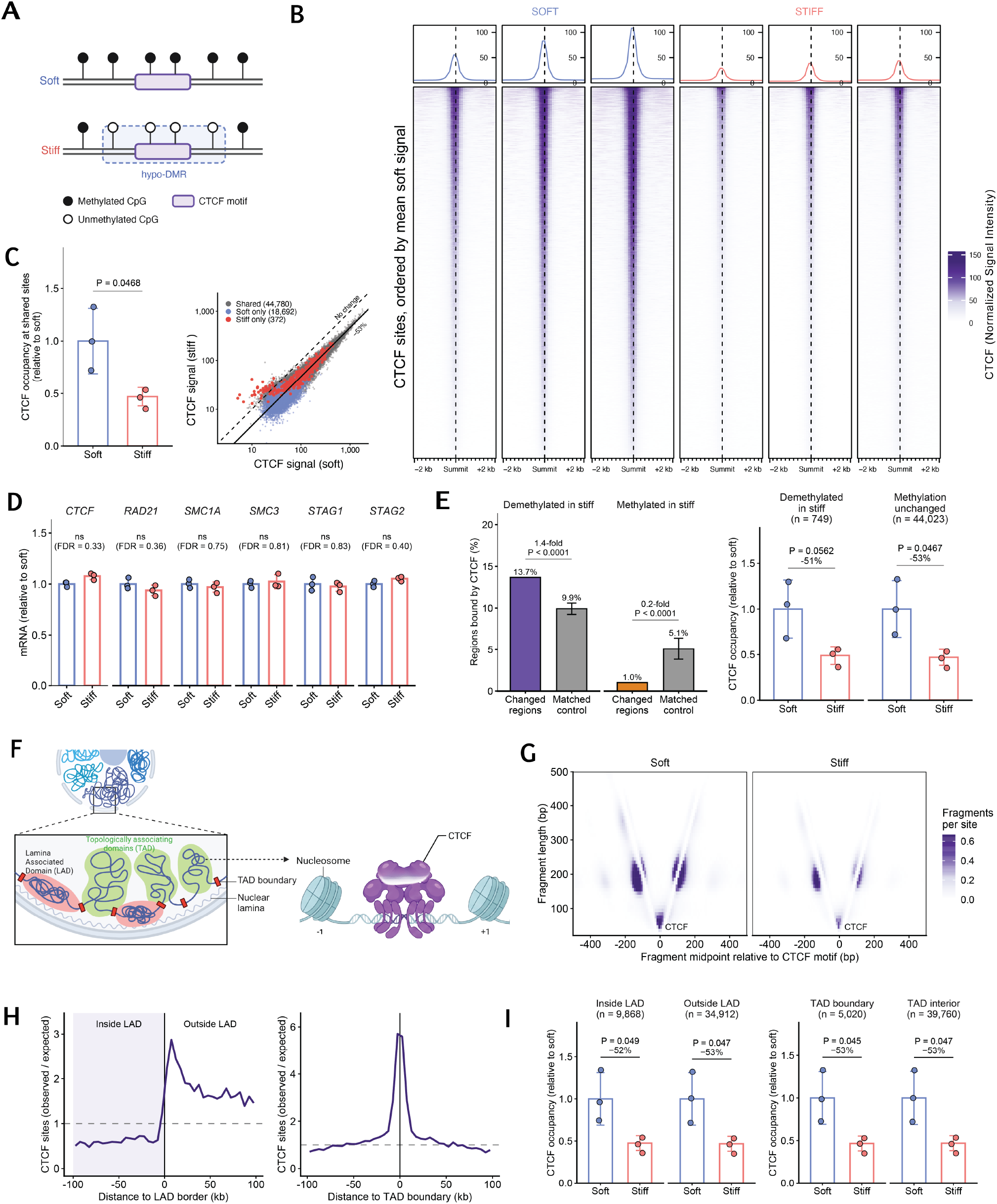
Matrix confinement reduces CTCF occupancy in GFs. **(A)** Schematic of CpG methylation at a CTCF motif in GFs in soft (G′ 0.75 kPa) and stiff (G′ 2.0 kPa) matrices. In the stiff condition, CTCF motifs lie within hypomethylated DMRs. **(B)** Heatmaps and mean profiles of CTCF CUT&RUN signal at 44,780 sites bound in both conditions (shared sites), ±2 kb around the CTCF summit, in GFs in soft and stiff hydrogels. Each column is one biological replicate (n = 3 per condition). Rows are ordered by mean soft signal. Signal was normalized to the yeast spike-in. **(C)** Left, total CTCF occupancy at shared sites relative to the soft mean. Data are presented as mean ± s.d.; dots represent biological replicates (n = 3). P-value determined by two-tailed unpaired t-test. Right, mean spike-in-normalized CTCF signal per site in soft and stiff matrices for all 63,844 CTCF sites. Colours indicate the condition in which a peak was detected. Dashed line, no change; solid line, overall stiff/soft ratio (0.47). 99.5% of sites lie below the no-change line. **(D)** mRNA expression of CTCF and cohesin subunits by bulk RNA sequencing (n = 3), relative to soft. Data are presented as mean ± s.d. FDR determined by DESeq2 Wald test with Benjamini–Hochberg correction. **(E)** Left, percentage of hypomethylated (demethylated in stiff) and hypermethylated (methylated in stiff) CpG DMRs that overlap a CTCF CUT&RUN site, compared with background regions matched for length, GC content and CpG density. Error bars indicate the 95% range of 10,000 matched background draws. P-values determined by empirical two-sided test against the matched background, for each DMR class separately. Right, CTCF occupancy at shared sites within hypo-DMRs (n = 749 sites) or outside DMRs (n = 44,023 sites), relative to soft. Data are presented as mean ± s.d. of biological replicates (n = 3). P-values determined by two-tailed unpaired t-test. **(F)** Schematic of lamina-associated domains (LADs), topologically associating domains (TADs) and TAD boundaries at the nuclear periphery. Inset, CTCF bound between flanking nucleosomes. **(G)** V-plots of CUT&RUN fragments at 24,266 motif-containing shared sites, oriented by motif strand. Fragment length is plotted against fragment midpoint position relative to the CTCF motif. Short fragments (≤120 bp) mark the CTCF footprint, and 150–260 bp fragments mark flanking nucleosomes. Mean of three biological replicates, spike-in normalized per site. **(H)** Density of shared CTCF sites relative to LAD borders (left) and TAD boundaries (right) defined in human HFFc6 fibroblasts, expressed as observed/expected relative to 500,000 random genomic positions in 5 kb bins. Dashed line, expected density. Negative distances in the left panel lie inside LADs. LADs, HFFc6 lamin B1 DamID domains present in both replicates (4DN Data Portal; 4DNFIN63IWZ3, 4DNFIHHIVIME). TAD boundaries, HFFc6 Micro-C insulation boundaries (4DN; 4DNFIC8438YF). **(I)** CTCF occupancy at shared sites located inside or outside HFFc6 LADs, and at or away from HFFc6 TAD boundaries, relative to soft. Data are presented as mean ± s.d.; dots represent biological replicates (n = 3). P-values determined by two-tailed unpaired t-test. Specific P-values are indicated on the graphs, where P < 0.05 was considered statistically significant. Panels A and F created with BioRender.com.

CTCF occupied 44,780 sites in both conditions (shared sites), distributed across promoters, introns, and distal intergenic regions (**Supplementary Fig. S16A**). CTCF occupancy at shared sites was lower in GFs in stiff-matrix, with reduced signal at 99.4% of shared sites (**Fig. 6B**). Total CTCF occupancy at shared sites was significantly reduced, by 53%, in the stiff condition (**Fig. 6C**). The reduction was proportional across all 63,844 CTCF sites, irrespective of site strength (**Fig. 6C**). CTCF and cohesin subunit (*RAD21, SMC1A, SMC3, STAG1, and STAG2*) transcript levels were unchanged between conditions (**Fig. 6D**), showing that reduced occupancy was independent of transcriptional regulation.

Hypo-DMRs in the stiff condition overlapped CTCF-bound sites more often than background regions matched for length, GC content, and CpG density (13.7% versus 9.9%), whereas hyper-DMRs were depleted of CTCF-bound sites (1.0% versus 5.1%) (**Fig. 6E**). This confirms that demethylation under confinement occurs preferentially at CTCF-bound sequences (**Fig. 5C**). However, CTCF sites within hypo-DMRs showed the same reduction in occupancy as sites outside DMRs (**Fig. 6E**), and methylation change did not correlate with CTCF occupancy change (Spearman ρ = −0.02; Supplementary Fig. S16C). Reduced CTCF occupancy under confinement is therefore independent of DNA methylation at the binding site. V-plot analysis of CUT&RUN fragment lengths^44^ showed CTCF centered on its motif between positioned flanking nucleosomes in both conditions, with lower signal in the stiff condition (**Fig. 6G**). CTCF remained positioned at its binding sites under confinement, with reduced occupancy. CTCF sites were mapped relative to LADs and TAD boundaries defined in human HFFc6 fibroblasts ^45,46^ (**Fig. 6F**). Shared CTCF sites were depleted within LADs and enriched at LAD borders and TAD boundaries (**Fig. 6H**). In the stiff condition, CTCF occupancy was reduced equally inside and outside LADs and at and away from TAD boundaries (**Fig. 6I**). Further, CTCF signal was reduced at promoter, exonic, intronic, 3′ UTR, and distal intergenic sites (**Supplementary Fig. S16B**).

These findings show that matrix stiffness-mediated confinement reduces CTCF occupancy uniformly at positioned binding sites, independent of CTCF and cohesin expression and local DNA methylation. A uniform reduction across all sites suggests a nucleus-wide effect of confinement rather than regulation at individual sites. Reduced CTCF occupancy is therefore a further feature of confinement-induced chromatin reorganization in GFs, together with H3K27Ac suppression, H3K9me3 redistribution, and methylation redistribution (**Figs. 3–5**). As CTCF marks domain boundaries, its reduced occupancy may contribute to the broader effective chromatin contact range in the confined state (**Fig. 5D–G**). Whether TAD boundaries and chromatin loops are altered in confined GFs remains to be determined by chromatin conformation capture assays.

### 7. Matrix stiffness governs differential epigenomic states at collagen synthesis and inflammatory gene loci in GFs

GFs maintain connective tissue integrity^3,8–11,47,48^ through ECM synthesis and innate immune surveillance^49^. We examined whether ECM stiffness-associated methylation differences translate to changes in collagen gene expression and TLR-mediated inflammatory responses. SHG imaging of healthy human gingival explants revealed organized fibrillar collagen networks oriented in stratified bands across tissue depth (**Fig. 7A, B**), confirming the architectural integrity of the stiff ECM matched by the stiff hydrogel condition^3^. GFs in stiff matrices showed transcriptional upregulation of multiple collagen genes, including COL5A3, COL4A2, COL6A5, and COL4A1 (**Fig. 7C**), consistent with the matrix-synthesizing phenotype maintained by stiff substrates in gingival fibroblasts^3,50^. Genome-wide DMR analysis showed that upregulated collagen genes carry promoter and gene body hypo-DMRs in the stiff condition (**Fig. 7D**), suggesting that targeted demethylation at these loci epigenetically licenses the matrix-synthesizing phenotype in GFs ^3,49–51^

**Figure 7.**
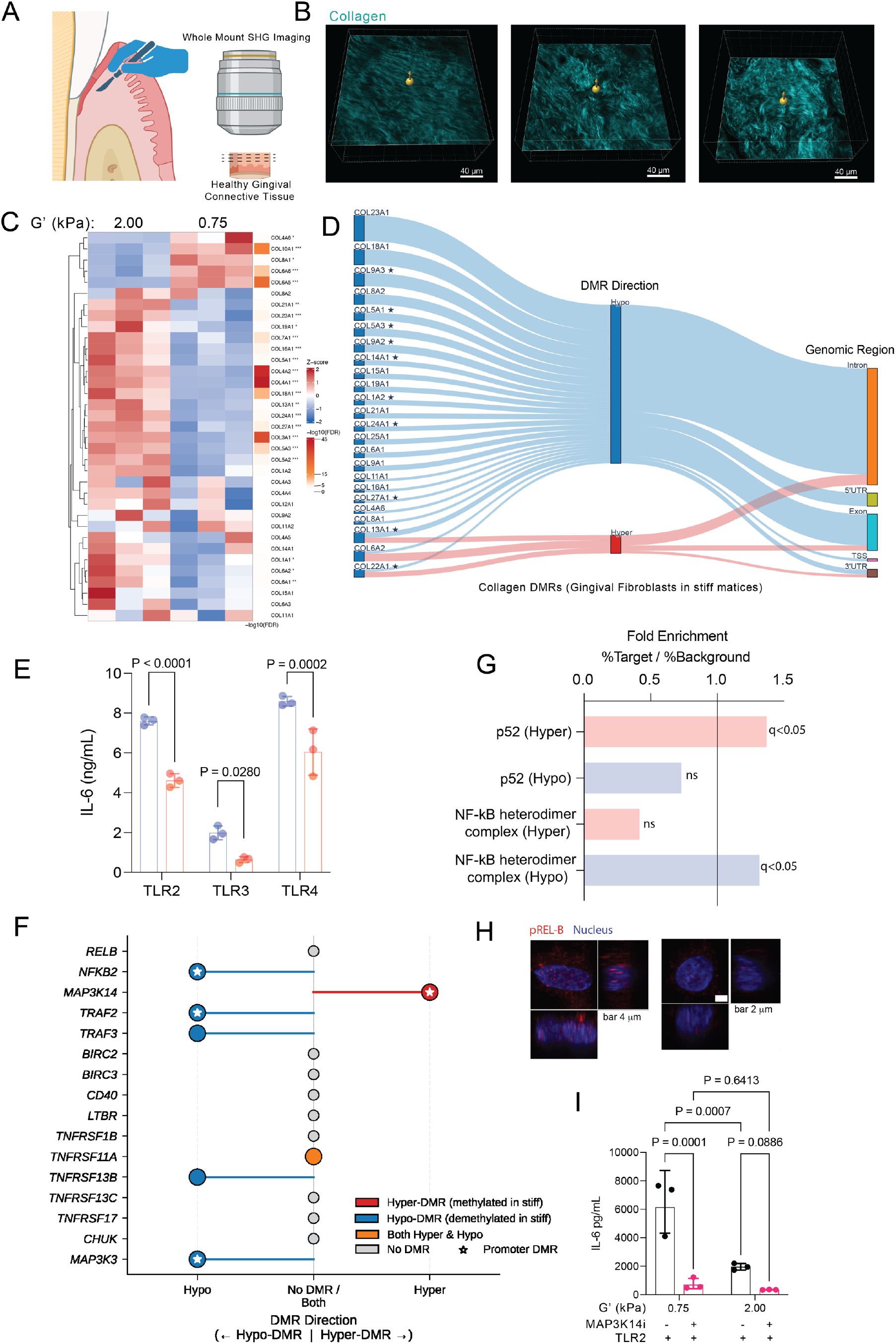
Confinement-driven methylation licenses the collagen synthesis program and silences non-canonical NF-κB inflammatory signaling through MAP3K14 promoter hypermethylation. (A) Schematic illustrating whole-mount SHG imaging workflow of healthy human gingival connective tissue. (B) Representative whole-mount SHG images showing fibrillar collagen architecture (cyan) in healthy human gingival connective tissue across three z-depth projections. Scale bars, 40 µm. (C) Heatmap showing z-scored expression of collagen genes in GFs in stiff (G’ 2.0 kPa) and soft (G’ 0.75 kPa) matrices from bulk RNA sequencing (n = 3). Color bar indicates −log₁₀(FDR). (D) Sankey diagram showing the direction of DMRs (hypo or hyper in the stiff condition) and their genomic region distribution for collagen genes in GFs in stiff matrices. Star symbols indicate promoter-localized DMRs. (E) ELISA quantification of IL-6 secretion from GFs in soft and stiff matrices following stimulation with TLR2, TLR3, and TLR4 agonists (n = 3). Data shown as mean ± s.d. P-values determined by two-way ANOVA with Tukey’s multiple comparisons test. (F) Lollipop plot showing the DMR status of non-canonical NF-κB pathway genes in GFs in stiff matrices. Hyper-DMR (red), Hypo-DMR (blue), both Hyper and Hypo (orange), no DMR (grey). Star symbols indicate promoter-localized DMRs. (G) Bar chart showing fold enrichment (%Target/%Background) of p52 and NF-κB heterodimer complex binding site motifs in hypermethylated (Hyper) and hypomethylated (Hypo) DMRs. q-values indicated. (H) Representative confocal immunofluorescence images of phosphorylated RelB (pREL-B, red) and nucleus (blue) in GFs in soft and stiff matrices. Scale bars, 4 µm and 2 µm. (I) ELISA quantification of IL-6 secretion from TLR2-activated GFs in soft (G’ 0.75 kPa) and stiff (G’ 2.0 kPa) matrices with or without MAP3K14 inhibitor (MAP3K14i) treatment (n = 3). Data shown as mean ± s.d. P-values determined by two-way ANOVA with Šídák’s multiple comparisons test. Panel A created with Biorendor.com

Beyond collagen synthesis, GFs function as innate immune sentinels through TLR signaling^3,52,53^, which recognizes bacterial lipoproteins, peptidoglycans, and lipoteichoic acid from the subgingival biofilm^53–55^. GFs in soft matrices produced significantly higher IL-6 following stimulation with TLR2, TLR3, and TLR4 (**Fig. 7E**). This shows that stiffness-dependent suppression of TLR-mediated IL-6 secretion is not receptor-specific but reflects a broader, mechanically regulated inflammatory setpoint. TLR2 was the focus of subsequent mechanistic analysis, given its established role as the primary pattern-recognition receptor in periodontal inflammation^49^. These findings extend our prior demonstration that stiff matrices suppress non-canonical NF-κB signaling and TLR-mediated cytokine secretion in GFs through actomyosin contractility and IKK activity^3^.

MAP3K14, encoding NIK, the initiating kinase of the non-canonical NF-κB pathway, carried a promoter-localized hyper-DMR in the stiff condition (**Fig. 6F**). Nuclear pREL-B signal was lower in stiff-matrix GFs (**Fig. 7H**), confirming attenuation of non-canonical pathway activation under confinement. Downstream pathway genes, including NFKB2 and TRAF2, carried promoter hypo-DMRs, showing that chromatin silencing is restricted to the MAP3K14 locus rather than distributed across the pathway. Without NIK kinase activity, downstream chromatin accessibility cannot drive pathway activation^56^. Further, p52 motifs were enriched within hypermethylated chromatin regions while NF-κB heterodimer motifs were enriched in hypomethylated regions (**Fig. 7G**). p52 homodimers lack a transactivation domain and displace transcriptionally activating p65/p50 heterodimers from NF-κB recognition sequences^56^, consistent with chromatin compaction and repressive NF-κB occupancy co-occurring at inflammatory loci in the stiff condition^3,50^. NIK inhibition in soft-matrix GFs reduced TLR2-stimulated IL-6 secretion to levels comparable with the stiff condition, whereas MAP3K14 inhibition in stiff-matrix cells produced no further reduction (Fig. 6I). MAP3K14 promoter hypermethylation produces the same suppression of non-canonical NF-κB-driven cytokine output as direct NIK inhibition. These findings establish matrix stiffness mediated confinement as a direct regulator of MAP3K14 promoter methylation and non-canonical NF-κB-dependent inflammatory output in stromal fibroblasts.^3,50^

### 8. Gingival fibroblasts retain a partial mechanical memory of prior matrix stiffness in the nuclear envelope

Stromal connective tissues undergo continuous ECM remodeling in health. Protease-mediated collagen degradation in inflammatory disease reduces matrix stiffness and removes the physical confinement that healthy ECM imposes on resident fibroblasts. Subsequent matrix deposition during repair restores confinement, exposing fibroblasts to cyclical transitions between soft and stiff mechanical environments. Whether the confinement-driven nuclear and chromatin changes described above reset upon return to a permissive matrix, or persist as a stiffness imprint independent of the current mechanical state, is unknown. To address this, GFs were primed in soft or stiff hydrogels for five days, retrieved by enzymatic digestion, and re-encapsulated into a second soft or stiff gel for 24 hours (**Fig. 8A**), generating four transfer conditions: soft-to-soft, soft-to-stiff, stiff-to-stiff, and stiff-to-soft.

**Figure 8.**
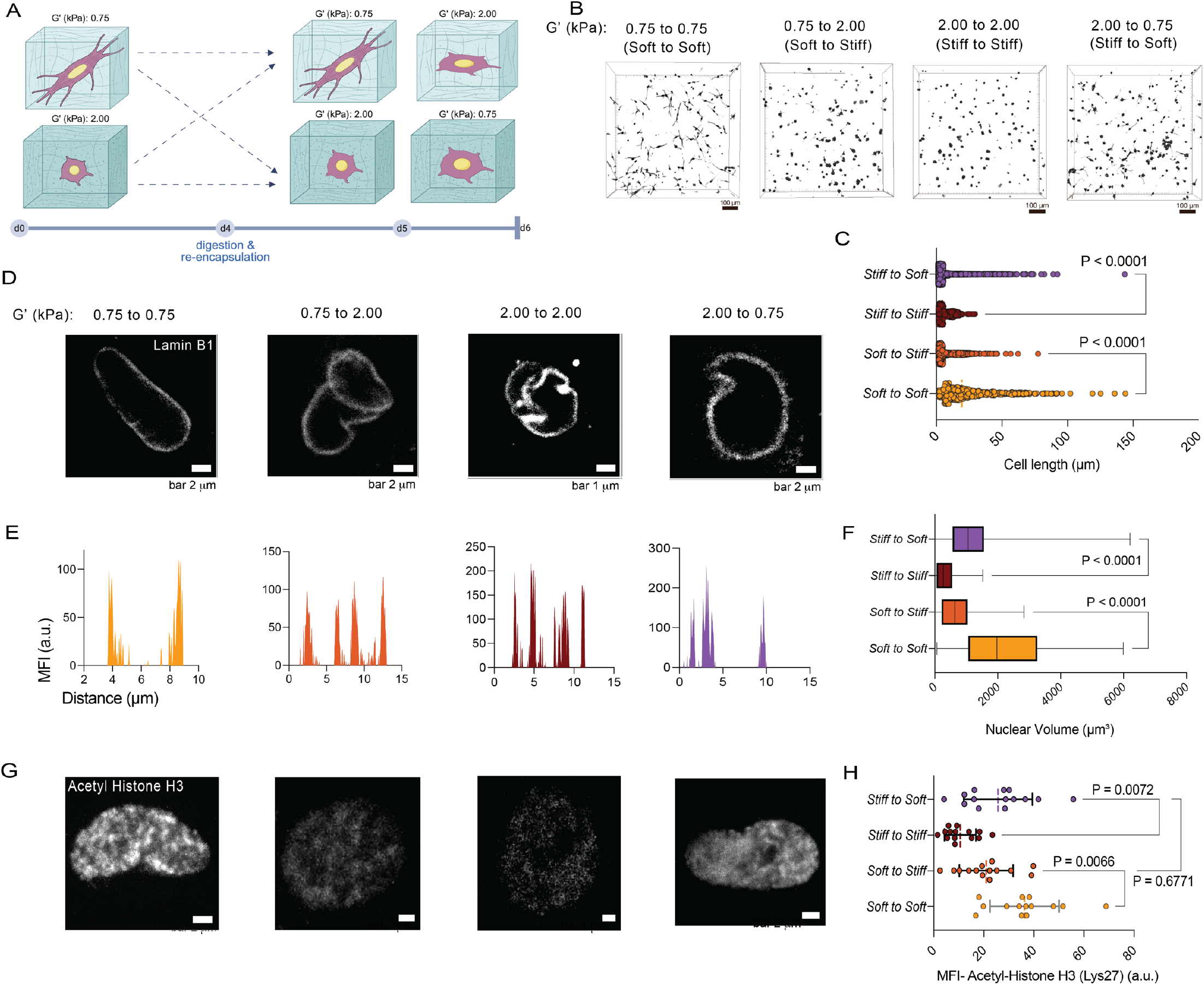
Gingival fibroblasts retain a mechanical memory of prior matrix stiffness. (A) Schematic of the mechanical memory experiment. GFs were primed in soft (G’ 0.75 kPa) or stiff (G’ 2.0 kPa) collagen-alginate hydrogels for five days, retrieved by enzymatic digestion, and re-encapsulated into a second soft or stiff gel for 24 hours, generating four transfer conditions (soft to soft, soft to stiff, stiff to stiff, stiff to soft). Created with BioRender.com. (B) Representative 3D confocal reconstructions of GF morphology in the four transfer conditions. Scale bars, 100 µm. (C) Quantification of cell length across transfer conditions. Dots represent individual cells pooled from three independent experiments. P-values were determined by two-way ANOVA followed by Šídák’s multiple comparisons test. (D) Representative Lamin B1 immunofluorescence images of GF nuclei (top) with corresponding intensity profile plot for each transfer condition. Two-peak profiles indicate smooth nuclear envelopes; multi-peak profiles indicate nuclear envelope folding. Scale bars as indicated. (E) Quantification of Acetyl-Histone H3 (Lys27) mean fluorescence intensity across transfer conditions and immunofluorescence images (right). Dots represent individual nuclei pooled from three independent experiments. P-values were determined by two-way ANOVA followed by Šídák’s multiple comparisons test. Data are presented as mean ± s.d., and specific P-values are indicated on the graphs, where P < 0.05 was considered statistically significant.

GFs re-encapsulated into soft gels spread and elongated, while cells re-encapsulated into stiff gels adopted a rounded morphology irrespective of prior priming. Cells primed in stiff and transferred to soft regained a spread morphology but remained shorter than cells primed and maintained in soft matrices (P < 0.0001), showing that cell morphology tracked the current matrix with a partial residual effect of mechanical history (**Fig. 8B, C**). Lamin B1 imaging revealed smooth nuclear envelopes in cells maintained continuously in soft matrices, whereas cells primed in stiff matrices retained folded nuclear envelopes after transfer to soft (**Fig. 8D**). Nuclear envelope folding persisted despite recovery of spread cell morphology, reflecting prior confinement independently of the current mechanical state. H3K27 acetylation showed greater reversibility. Cells primed in stiff and transferred to soft recovered acetylation to levels indistinguishable from soft-maintained cells, showing that the current matrix resets the chromatin acetylation state (**Fig. 8E**). Cells primed in soft and transferred to stiff retained higher acetylation than stiff-maintained cells while falling below soft-maintained levels, showing that a single stiff transfer did not fully erase soft priming. Nuclear envelope architecture therefore retained mechanical history more persistently than H3K27 acetylation. These results show that stiffness-driven chromatin changes in fibroblasts are largely reversible^57,58^, whereas nuclear envelope folding partially persists after matrix softening^6,59^.

## Conclusion, Limitations, and Future Outlook

This study shows that ECM stiffness regulates the chromatin and inflammatory state of gingival fibroblasts through mechanical confinement. In stiff matrices, confinement compacts the nucleus and folds the nuclear envelope independent of actomyosin contractility. The confined nucleus in stiff matrix maintains a distinct chromatin state, with suppressed H3K27Ac, redistributed H3K9me3, methylation gain at pericentromeric satellite repeats, hypomethylation at collagen-synthesis gene promoters and CTCF-binding motifs, and reduced CTCF occupancy across the genome. This state extends to inflammatory signaling, as MAP3K14 promoter hypermethylation attenuates non-canonical NF-κB activity and limits TLR-induced IL-6 secretion. Reduced confinement in soft matrices relieves this chromatin state and renders fibroblasts susceptible to inflammatory activation, while nuclear envelope folding is partially retained as a mechanical memory of prior confinement (Fig. 9).

**Figure 9.**
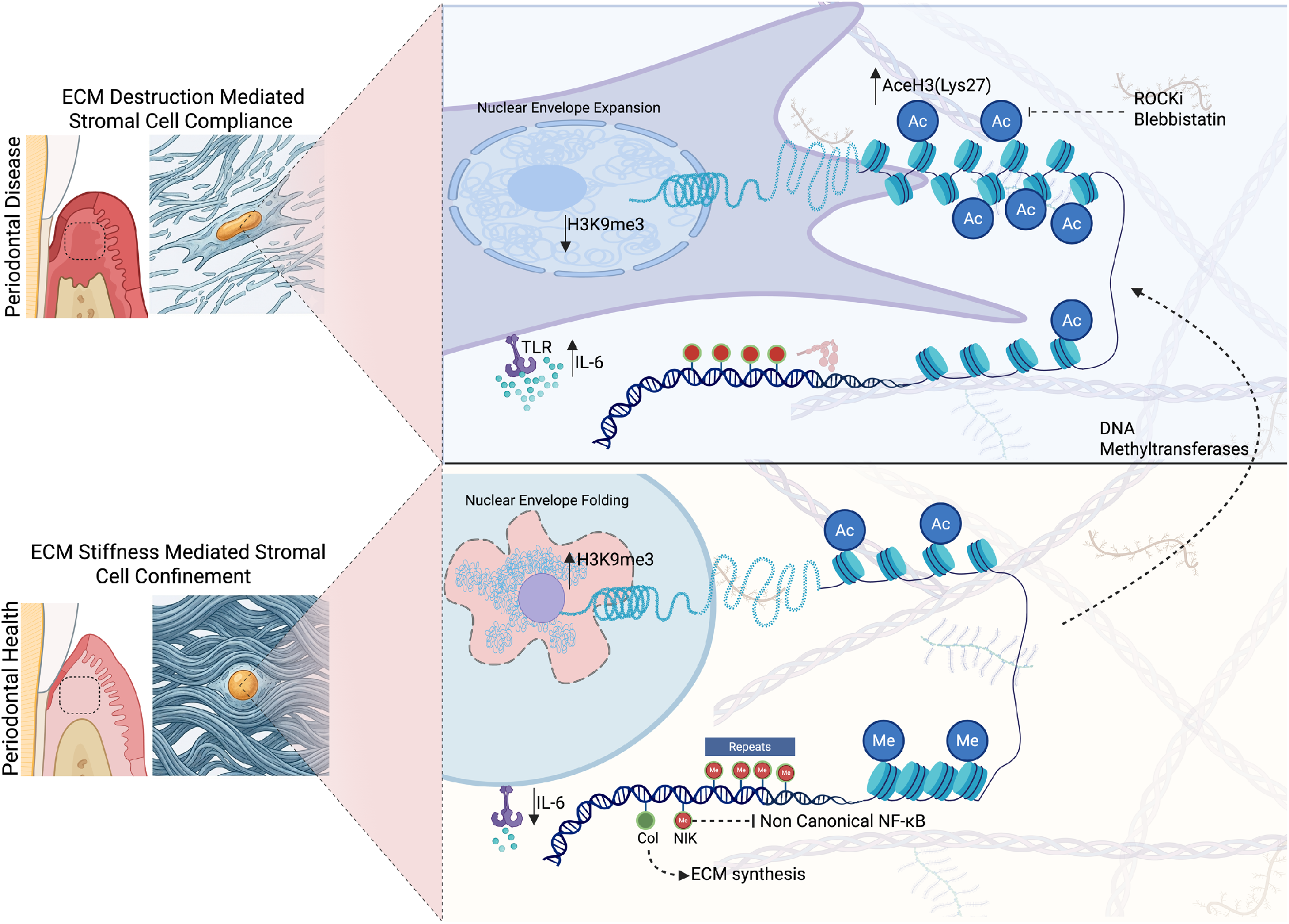
Proposed model for ECM stiffness-dependent epigenomic regulation of gingival fibroblast identity in periodontal health and disease. Created with Biorendor.com.

ECM degradation is a shared pathological process in chronic inflammatory connective tissue diseases, including periodontitis and musculoskeletal and gastrointestinal inflammatory diseases. Protease-mediated matrix degradation softens the tissue and reduces the physical confinement imposed by the intact matrix on resident fibroblasts. Our findings show that reduced confinement shifts fibroblasts from a low-acetylation to an inflammation-permissive chromatin state. Matrix softening is therefore not only a consequence of inflammation but may also be a mechanical driver of the fibroblast inflammatory state. Using donor-derived human gingival fibroblasts in hydrogels in which stiffness is decoupled from ligand density and polymer architecture, together with patient-derived tissue biopsies, this work isolates mechanical confinement as the causal variable in primary human cells and links it to the human tissue state. The confinement-regulated chromatin state described here may represent a conserved stromal mechanism across tissues, and restoring matrix mechanics may offer a route to restoring stromal homeostasis^6^.

There are several limitations of these studies. Whether reduced CTCF occupancy alters TAD boundaries and chromatin loops in confined GFs remains to be determined by chromatin conformation capture assays. LADs and TAD boundaries were defined from reference fibroblast maps rather than from GFs. The DNMT inhibitor experiment cannot separate the contribution of methylation loss from the accompanying morphological changes during H3K27Ac restoration, a distinction that selective DNMT1 knockdown would resolve. At the tissue level, the data are correlational across a heterogeneous stromal compartment, and the therapeutic value of MAP3K14 remains to be established in vivo. Future studies will examine whether restoring ECM stiffness reverses the confinement-associated chromatin state, including CTCF occupancy, and whether MAP3K14 can be targeted to restore stromal immune homeostasis in inflammatory disease.

## Ethical Statement and Human Tissue Procurement

All human gingival tissues and donor-derived gingival fibroblasts (GFs) utilized in this research were obtained from the University of Pennsylvania Periodontology Clinic under Institutional Review Board oversight (IRB #844933; PI: Ko). These materials, which included deidentified human gingival biopsies from healthy donors, were used to establish ex vivo explant models and isolate primary GFs for 3D hydrogel systems. To maintain donor privacy, all human tissue samples were deidentified before being processed for experimental use. Furthermore, written informed consent was obtained from all participants prior to the collection and use of human tissues in accordance with ethical standards and institutional protocols.

## Acknowledgements

This work was supported by the National Institute of Dental and Craniofacial Research (NIDCR) through a training grant to the Center for Innovation & Precision Dentistry (CiPD) (R90DE031532 to Hardik Makkar) and partially by NIDCR Award R01DE035592 (KK). Additional support was provided in part by the Collaborative Research Grant from the Institute for Regenerative Medicine in the Perelman School of Medicine and the School of Dental Medicine at the University of Pennsylvania (Vining). This study was also partially supported by the National Institute of General Medical Sciences (NIGMS) (R35GM157079, Vining), by NCI Award U54CA261694 (VV, V.B.S.), NSF CEMB Grant CMMI-154857 (VV, R.G.W., V.B.S.), NSF Grant DMS-2347834 (VV, V.B.S.), NIBIB Awards R01EB017753 (VV, V.B.S) and R01EB030876 (VV, V.B.S.), and NIGMS Award R01GM155943 (VV, V.B.S). We thank Gordon Ruthel at the Penn Vet Imaging Core for assistance with confocal microscopy and SHG imaging. Confocal microscopy was performed on an instrument purchased with support from an NIH Shared Instrumentation Grant (S10 OD032305-01A1). Mobilized CD34+ HSPCs were obtained from the Fred Hutchinson Cancer Center Cooperative Centers of Excellence in Hematology (CCEH) Cell Procurement & Processing Core, which is supported by U54 DK106829. We thank Kawin Prasongyuenyong from Penn Dental Medicine (Ko Lab) and Edgardo Arroyo at the Penn Center for Musculoskeletal Disorders histology core for assistance in tissue processing. Histology core service was performed with support from grant P30 AR069619.

## Competing Interest

The authors declare no competing interests.

## Data Availability

All data will be available from a publicly posted Dryad repository upon acceptance of the article at a DOI to be provided. Sequencing data will be deposited on the GEO database.

## Notes

### Competing Interest Statement

The authors have declared no competing interest.

### Summary of Updates

Updated manuscript, supplement, authors, funding, and abstract

