## Supplementary Information for "Nuclear confinement from matrix stiffness drives epigenomic reprogramming of gingival fibroblasts"

Hardik Makkar 0000-0002-8416-4138

Vinayak Vinayak 0000-0002-4597-5634

Kang I Ko 0000-0003-2967-8846

Rebecca G. Wells 0000-0002-6988-4102

Vivek B. Shenoy 0000-0002-2645-1016

Kyle H. Vining 0000-0002-4009-879X

**Methods**

**Isolation of Donor-Derived Gingival Fibroblasts**

Gingival fibroblasts were isolated from healthy human gingival tissue from the University of Pennsylvania, Periodontology Clinic (IRB exemption #844933, PI: Ko). Freshly obtained gingival tissues were washed in 1x HBSS without calcium and magnesium supplemented with antibiotic-antimycotic (1X, Gibco). The gingival tissues were cut into small pieces and placed in a T25 flask for 72 hours undisturbed and supplemented with Dulbecco’s modified eagles medium (low glucose;1g/L) with 10% FBS and antibiotic-antimycotic (1X, Gibco). Cells were expanded, cryopreserved, and passages 2-4 were used for experiments.

**Preparation of Gingival ECM hydrogels**

Gingival ECM hydrogels were prepared as described previously^1-4^. Briefly, alginate (Pronova) was dissolved in a solution of 1x HBSS without calcium nor magnesium (14175095, Gibco) with HBSS/HEPES buffer) to a final w/v concentration of 5%. On the day of IPN fabrication, in a glass vial with a magnetic stir bar, bovine telo collagen I (Advanced Biomatrix) was neutralized on ice in v/v ratio of 17:3 with a solution created from 10x HEPES (14180046, ThermoFisher), 1 M HEPES, 1x HBSS, and 1M sodium hydroxide (NaOH) mixed in a v/v ratio of 10:2:2:1, respectively. Extra NaOH was added dropwise until the phenol red indicated a pH of ~7. HBSS/HEPES buffer was used to adjust the final concentration of collagen to 4 mg/mL. Subsequently, a 100 mg/mL CaCO3 slurry was prepared in endotoxin-free ultra-pure water. With these solutions ready, an aliquot of the CaCO3 slurry was incorporated into the neutralized collagen at a final concentration of 10, 20, or 30 mM. Then, alginate was added to the mixture at a final w/v concentration of 1%. These solutions were mixed with a magnetic stirrer. Next, a solution of 0.4 g/mL glucono-delta-lactone was formulated with HBSS/HEPES buffer immediately before its incorporation into the gel mixture at a 4x molar excess of CaCO3. The solution was stirred and mixed by pipetting. Working quickly, the liquid hydrogel solution was casted in 96 (60 µL) or 48 well-plates (150 µL) and incubated at 37°C for gelation and further experimentation.

**Rheological Characterization of hydrogels and gingival tissues**

The rheological properties of the gingival ECM hydrogels were measured on the HR-30 rheometer (TA Instrument) as shown previously^4^. 100 µL of the hydrogel solution was dropped on top of the 20 mm Peltier plate at 37°C and loaded with a 20 mm parallel plate geometry. Excess solution was wiped off, and a solvent trap sealed with mineral oil was added to prevent dehydration. Time sweep was conducted at an oscillation of 1 Hz and strain of 2% until both storage modulus (G’) and loss tangent (tan δ) reached an equilibrium (1 to 2 h), after which a frequency sweep from 0.1 to 10 Hz was performed under 2% strain.

**Cell culture and hydrogel encapsulation**

To prevent collagen contraction, surface modification of 96 and 48 flat-bottom culture plates was done with a 2 mg/ml dopamine solution (Sigma Aldrich, USA) prepared in 10 mM Tris-HCl (pH 8.5) for 2h at room temperature^5^. After 2 hours of incubation, the dopamine solution was gently aspirated, and the surfaces were washed twice with sterile filtered deionized (DI) water to remove unbound dopamine. The washed substrates were kept sterile at room temperature until use. Gingival Fibroblasts (GFs) were isolated from healthy donor gingival tissues, cultured as explants. The isolated cells were cultured in Dulbecco’s modified Eagle's medium (low glucose;1g/L) with 10% FBS. GFs suspended in HBSS/HEPES buffer and added to the collagen mixture, prior to the addition of alginate, at a final concentration of 1E06 cells/mL. Once all the hydrogel components were added and mixed, 60 µL or 150 µL hydrogel constructs were prepared in a 96- or 48-well flat-bottom well plates. The plates were then moved to a cell incubator at 37°C and 5% CO2. After 30 minutes, HBSS/HEPES buffer was added to the hydrogels, to maintain a pH ~7. Later, the buffer in the wells containing the ECM hydrogels was replaced with a low-serum defined media^6^.

**Exposure of gingival ECM hydrogels to pattern recognition receptor agonists**

Gingival ECM hydrogels cultures for four days were exposed with TLR-2 agonist (Pam3CSK4, Invivogen), TLR-4 agonist (ultrapure *E. coli* LPS, Invivogen), TLR-3 agonist for 24 h (Supplementary table T2). Following challenge, the culture supernatants were collected and stored at −80°C for downstream cytokine analysis.

**Retrieval of cells from ECM hydrogels and Viability assessment**

Encapsulated cells were recovered by enzymatic digestion of the hydrogels^1-3,7^. Briefly, hydrogels were incubated with a digestion solution containing 300 U/mL collagenase type I (Thermo Scientific) and 34 U/mL alginate lyase (Sigma-Aldrich) in Dulbecco’s phosphate-buffered saline (DPBS) supplemented with calcium and magnesium, and 0.5% bovine serum albumin (BSA), at 37°C for 15 minutes. An additional 60–150 µL of 300 U/mL collagenase I was then added with a second incubation at 37°C for 40 minutes. To terminate digestion, 0.5 mL of MACS buffer (DPBS without calcium and magnesium, supplemented with 2 mM EDTA and 0.5% BSA) was added, and the wells were washed using ice-cold PBS -/- and EasySep buffer (Stem Cell Technologies). Additional washes were performed if cells remain attached, or if undigested hydrogel was visible. The cell suspension was centrifuged at 400 × g for 5 minutes at 4°C for three times to obtain a clean cell pellet for downstream experiments.

**Enzyme Linked Immunosorbent Assay (ELISA) for secretome analysis**

In accordance with manufacturer’s instructions, ELISAs for cytokines IL-6 (Biolegend) were performed using the culture supernatants. The absolute cytokine values were normalized to the total protein content using BCA assay (Thermo Fisher Scientific).

**Second Harmonic Generation Imaging and image analysis.**

Second harmonic generation (SHG) imaging of formalin fixed paraffin embedded sections of human gingiva and viable whole gingival tissues were collected on Leica SP8-MP  Upright with an external non-descanned hybrid detector (HyD – RLD2). The laser was tuned to  910nm, and the filter associated with the lens was 435~485nm. Confocal imaging was performed on a Leica Stellaris DMi8 microscope under appropriate fluorescence imaging settings for the respective samples. Z-stacks were acquired and processed in FIJI (ImageJ) and Imaris software (Oxford Instruments) using both forward and backward SHG signals.

**Immunostaining**

The Gingival ECM hydrogels were fixed in 4% PFA (EMS), processed, and embedded in optimum cutting temperature compound (Tissue-Tek). Tissue sections (10 µm thick). For immunostaining^8^, the sections were first washed with DPBS, antigen retrieval with 0.5% Triton-X followed by blocking of nonspecific staining. The sections were incubated with respective primary antibodies overnight at 4°C, washed and incubated with respective secondary antibodies for 2 hours at 4°C (Supplementary table T3). The nuclei were counterstained with DAPI and mounted with anti-fade fluorescent mounting medium (Abcam). The slides were visualized under confocal microscope (Leica Stellaris, Leica Microsystems equipped with Leica Application Suite X software). Whole mount immunostaining was done as previously described^9^. Briefly, antigen retrieval and permeabilization was done by immersing the hydrogels in 0.5% Triton-X in phosphate-buffered saline (PBS) under orbital shaking for 2 h. The gels were then incubated in blocking solution (PBS containing 0.5% Triton-X, 10% goat serum, and 5% bovine serum albumin) overnight under orbital shaking. Tissues were incubated with primary antibodies (Supplementary table T3) for 2 d at 4 °C followed by washing under orbital shaking for 3 h. This was followed by incubation with respective secondary antibodies (Supplementary table T3) for 2 d at 4 °C and counterstaining with DAPI. Whole mounts were visualized using laser scanning confocal microscopy (Leica Stellaris, Leica Microsystems). Image analysis was done using Fiji (NIH) and Imaris software (Oxford Instruments).

For immunofluorescence staining of formalin-fixed paraffin-embedded gingival biopsies, the tissues were fixed in 4% PFA (Sigma-Aldrich), processed, embedded in paraffin and sectioned (5 µm thick). Immunostaining of the deparaffinized tissue sections was performed by subjecting them to heat-induced epitope recovery at 121^o^C in a pressure vessel (Retriever™2100, Aptum Biologics) and 0.01M (pH6) citrate buffer (EMS). The sections were blocked of non-specific staining and incubated with respective primary antibodies (Supplementary Table T2) overnight at 4oC. This was followed by washing, labeling with appropriate secondary antibodies (Supplementary Table T2), counterstaining with DAPI, and mounting with anti-fade fluorescent mounting medium (Abcam). Immunostained slides were imaged using Leica SP8-MP microscope (Leica DMi8, Leica Microsystems equipped with Leica Application Suite X software)

**Bulk RNA- sequencing**

Cells encapsulated in hydrogels were harvested and lysed using RLT buffer supplemented with 1% β-mercaptoethanol (β-ME) (Sigma-Aldrich), following standard protocols for disruption and homogenization of cell pellets. Briefly, cell pellets were resuspended in 350 µL of RLT + β-ME and pipetted vigorously to ensure complete lysis and homogenization. Lysates were stored at –80°C or processed immediately for RNA extraction. Total RNA was isolated using a commercial column-based purification kit (RNeasy Mini Kit, Qiagen), according to the manufacturer’s instructions. This included an on-column DNase I digestion step to remove genomic DNA contamination and elution in RNase-free water. RNA concentration and integrity were assessed by spectrophotometry (NanoDrop) prior to sequencing. Bulk RNA sequencing was performed by Azenta Life Sciences. Library preparation, quality control, sequencing, and initial data processing were conducted by Azenta.

**Whole Genome Bisulfite Sequencing**

Genomic DNA was isolated from human gingival fibroblast samples using standard silica-column or magnetic-bead-based extraction protocols, and its concentration and purity were assessed spectrophotometrically via NanoDrop (Thermo Fisher Scientific) and fluorometrically using a Qubit dsDNA BR Assay Kit (Thermo Fisher Scientific). To establish an internal control for quantifying downstream bisulfite conversion efficiency, 1 µg of purified genomic DNA was spiked with 0.5% (w/w) unmethylated lambda phage DNA (Promega) before being physically fragmented via focused acoustic shearing using a Covaris S220 ultrasonicator (Covaris, Inc.) to isolate fragments distributed within a 200–400 base pair (bp) target range. Fragmented double-stranded DNA underwent enzymatic end-repair to yield blunt ends, followed by the addition of a single 3′ adenine (A-tailing) using an automated library preparation suite to facilitate the ligation of full-length, methylated barcoded adapters synthesized with 5-methylcytosine (5mC) at all cytosine positions, thereby protecting adapter-specific primer-binding sites from deamination during chemical treatment. Chemical bisulfite conversion was executed using the EZ DNA Methylation-Gold Kit (Zymo Research) to induce hydrolytic deamination of unmethylated cytosine residues into uracil while leaving protected 5mC and 5-hydroxymethylcytosine (5hmC) residues unreactive. The resulting single-stranded, bisulfite-converted fragments were purified and size-selected using Agencourt AMPure XP magnetic beads (Beckman Coulter) to eliminate adapter dimers before undergoing polymerase chain reaction (PCR) amplification with KAPA HiFi HotStart Uracil+ ReadyMix (2x) (Kapa Biosystems), an engineered formulation that bypasses template uracil bases without polymerase stalling. Final library constructs were quantified via a Qubit 2.0 Fluorometer (Life Technologies), verified by quantitative PCR (qPCR) to determine absolute molar concentrations of clonally amplifiable molecules, and evaluated for size distribution and structural integrity using an Agilent 2100 Bioanalyzer (Agilent Technologies). Validated libraries were pooled in equimolar ratios and subjected to high-depth whole-genome bisulfite sequencing (WGBS) on an Illumina NovaSeq 6000 system using a 150-bp paired-end (PE150) strategy. The raw FASTQ sequencing reads were preprocessed using FastQC (v0.11.9) to evaluate base-calling quality, and artifact removal, adapter trimming, and quality filtering were executed via Trim Galore! (v0.6.6) . Data filtering criteria strictly enforced the complete elimination of adapter-contaminated reads, the removal of reads where indeterminate bases (N) exceeded 10% of the total read length, and the exclusion of low-quality sequences where more than 50% of the constituent bases exhibited a Phred quality score (Q-score) less than or equal to 5 or individual bases presented a Q-score less than 20 (discarding truncated reads under 20 bp), ultimately yielding high-quality clean reads for downstream genomic alignment and differential methylation analysis.

**Differential Methylation Analysis**

Differentially methylated regions (DMRs) between stiff and soft conditions were identified genome-wide using a sliding-window approach. Regions meeting both statistical significance and effect-size thresholds were retained. Each DMR was annotated by chromosomal coordinates (0-based, GRCh38), the number of covered CpG sites (nCG), per-condition mean β values, and the signed methylation difference (Δβ = βstiff − βsoft). Regions with Δβ > 0 were classified as hyper-DMRs, and regions with Δβ < 0 were classified as hypo-DMRs. Hyper-DMRs therefore represent gain of CpG methylation under stiff substrate conditions. The area statistic (areaStat = Σ|Δβi| across all nCG sites within a region; percentage units, 0–100 scale) was used as an integrated summary of DMR effect size, capturing both per-CpG methylation magnitude and affected genomic span. Only CG-context cytosines are reported.

DMRs were annotated by intersection with GENCODE gene models and RepeatMasker elements (GRCh38). When a DMR overlapped multiple genomic features, a single primary annotation was assigned according to the following priority: promoter (TSS ± 2 kb) > CpG island > exon > 5′ UTR > 3′ UTR > CpG island shore > intron > repeat element > intergenic. All overlapping feature labels and gene symbols were retained as semicolon-delimited fields. Records with identical genomic coordinates were deduplicated to one entry per unique region. After deduplication, chromosome 19 contained 72 hyper-DMRs and 888 hypo-DMRs (960 regions total), corresponding to a hyper-DMR density of 1.23 DMRs per Mb. Regions within each directional class were ranked by descending |areaStat|, with rank 1 indicating the largest effect.

Chromosome 19 was divided into 18 non-overlapping bins by the WGBS analysis pipeline. Bins 1–17 were each 3,276,800 bp, and the terminal bin spanned the remaining 2,912,016 bp. Three density tracks were extracted for each bin: CG methylation density, gene density and transposable element (TE) coverage. CG methylation density was calculated as the mean β value across all covered CpG sites per bin and per sample; binned ΔCG methylation was expressed as (βstiff − βsoft) × 100. Gene density was defined as the number of GENCODE protein-coding gene transcription start sites (TSSs) within each bin. TE coverage was defined as the fraction of each bin overlapping RepeatMasker-annotated repetitive elements, including LINEs, SINEs, DNA transposons and LTR retrotransposons, and is reported as a percentage. The chromosome 19 mean TE coverage was 56.85%, calculated as the unweighted mean across all 18 bins. For visualization, a four-track ideogram was assembled in R using ggplot2, ggrepel and patchwork. Binned ΔCG methylation was displayed as per-bin bars centred at the bin midpoint and spanning 90% of the bin interval, with hypermethylation shown in orange and hypomethylation shown in blue.

### **Computational modelling of confinement-driven DNA methylation redistribution**

**Model construction and SOFT reference landscape-** To test whether confinement-driven changes in chromatin contacts are sufficient to redistribute DNA methylation, we developed a coarse-grained stochastic model of a heterogeneous methylation landscape. The model is defined on a one-dimensional genomic lattice and does not explicitly represent genes, repeat families, transcription-factor binding sites, topologically associating domains, or chromosome-specific annotations. Instead, we ask how an established SOFT methylation architecture responds when only the effective chromatin-contact landscape is changed.^10-12^

We represent the model chromosome as N = 1,200 consecutive 10-kb bins, corresponding to a 12-Mb genomic region. Each bin i carries a binary methylation variable s_i ∈ {−1,+1}, where +1 and −1 denote the methylated and unmethylated model states, respectively. Simulations are performed as stochastic ensembles. The methylation level at bin i is therefore defined as m_i = P(s_i = +1), estimated as the fraction of stochastic realizations in the +1 state. All model comparisons are made using this ensemble methylation profile rather than an individual binary realization.

The starting SOFT landscape was constrained by the experimental 10-kb WGBS architecture. SOFT bins were classified as high methylation (m ≥ 0.60), intermediate methylation (0.20 < m < 0.60), or low methylation (m ≤ 0.20). The corresponding experimental fractions were 0.67809, 0.31963, and 0.00228. We also used the experimentally measured run-length distributions of these states: high-methylation runs had a mean length of 65.5 kb and a median of 20 kb, intermediate runs had a mean of 30.8 kb and a median of 10 kb, and low-methylation runs had a mean of 11.7 kb and a median of 10 kb.

To generate an empirical-like SOFT track, the probability of starting a run in state q was taken proportional to f_q/L_q, where f_q is the observed genomic fraction of that state and L_q is its observed mean run length. Run lengths were drawn from lognormal distributions defined by the observed mean and median of each state and rounded to integer 10-kb bins. Consecutive runs were constrained to have different labels so that adjacent identical draws did not merge into an unintended longer run. Within each run, methylation values were sampled within the corresponding state range to avoid a piecewise-constant reference profile. The final SOFT reference had a global mean methylation of approximately 0.680. No STIFF methylation measurement was used to construct this reference.

**Fixed material classes and local fields-** Each bin was assigned a fixed coarse material label q_i from the SOFT reference. Bins with m_i ≥ 0.60 were assigned H-like material, whereas bins with m_i < 0.60 were assigned A-like material. These labels remain fixed during a simulation; only s_i evolves. We use H-like and A-like strictly as model material classes derived from the baseline SOFT methylation architecture. They are not experimentally annotated heterochromatin/euchromatin, repeat/regulatory compartments, CTCF domains, or TADs.

The two material classes were assigned centered local fields h_H = +0.23428517 and h_A = −0.23428517, giving Δh = 0.46857034. A common offset μ = −0.19635939 was then applied, giving total single-bin fields b_H = h_H + μ = +0.03792578 and b_A = h_A + μ = −0.43064456. These parameters were determined from the SOFT reference and frozen before the STIFF branch was evaluated. We did not use STIFF outcomes, locus-specific experimental responses, or genomic annotations to determine the fields.

#### Effective contacts and energetic model

The methylation configuration is governed by a material-selective contact Hamiltonian,

$$E_{\lambda}\left( \mathbf{s} \right)=-J\sum_{i<j} c_{ij}W\left( \left| i-j \right|;\lambda\right)s_{i}s_{j}-\sum_{i} \left( h_{i}+\mu\right)s_{i}.$$

Here, J = 0.8 is the cooperative coupling strength. Material selectivity is introduced through c_ij. The primary model uses η_cross = 0, so cooperative reinforcement occurs only between bins of the same coarse material class. This choice does not prohibit exchange between H-like and A-like bins; it only removes cross-material energetic reinforcement. Non-zero cross-material coupling is examined separately in the robustness analysis.

$$c_{ij}=\left\{ \begin{matrix} 1, & q_{i}=q_{j}, \\ \eta_{\mathrm{cross}}, & q_{i}\neq q_{j}. \end{matrix} \right.$$

Effective chromatin contacts are represented by a Gaussian kernel over genomic separation d,

$$W\left( d;\lambda\right)=\frac{\exp\left[ -d^{2}/\left( 2\lambda^{2} \right) \right]}{Z_{\mathrm{SOFT}}}, 1\leq d\leq4\lambda.$$

$$Z_{\mathrm{SOFT}}=\sum_{d=1}^{8} \exp\left[ -\frac{d^{2}}{2\times2^{2}} \right]=2.006584.$$

The same SOFT normalization factor is used in both conditions. This is important because increasing λ then changes both the genomic range and the integrated weight of effective contacts. For SOFT, λ = 2 gives an 8-bin cutoff (80 kb) and a one-sided integrated contact weight of 1.00. For STIFF, λ = 5 gives a 20-bin cutoff (200 kb) and an integrated weight of 2.873699. Thus, confinement is represented by a broader and denser effective contact landscape, while all other model parameters remain unchanged. λ is a reduced genomic interaction-range parameter that summarizes the altered 3D chromatin-contact environment; it is not a diffusion length for methyl groups.

For a bin i, the instantaneous local field is

$$F_{i}=h_{i}+\mu+J\sum_{j} c_{ij}W\left( \left| i-j \right|;\lambda\right)s_{j}.$$

where the sum includes bins within the truncated kernel support on either side of i, subject to the modeled chromosome boundaries.

#### Stochastic dynamics

**Contact-mediated exchange-** We use a Monte Carlo exchange process to represent contact-mediated redistribution of epigenetic state^13,14^. One Monte Carlo step (MCS) contains N exchange proposals, corresponding on average to one proposal per genomic bin. For each proposal, a source bin i is selected uniformly. A genomic separation d is drawn from the condition-specific contact kernel after normalizing the kernel weights to a partner-selection probability, and the left or right direction is chosen with equal probability. Proposals that fall outside the modeled chromosome are rejected rather than renormalized at the edge. If the selected bins have the same methylation state, no change occurs. If the states differ, they are exchanged and the move is accepted with the Metropolis criterion

$$P_{\mathrm{accept}}=min\left[ 1,exp\left( -\beta\Delta E \right) \right], \beta=1.0.$$

where ΔE is the exact change in the material-selective Hamiltonian. We independently compared the local ΔE expression used during simulation with the full Hamiltonian difference before and after a swap; the two agreed to numerical precision. Because exchange conserves the number of +1 and −1 states within each realization, it represents redistribution of epigenetic state rather than literal movement of methyl groups along the genome.

**Background turnover-** A second, rare process allows local methylation gain or loss. For a simulation of duration t MCS, the number of turnover attempts is drawn independently from a Poisson distribution

$$N_{\mathrm{rxn}}\sim Poisson\left( \gamma Nt \right), \gamma=0.001 \mathrm{bin}^{-1} \mathrm{MCS}^{-1}.$$

For each turnover event, a bin is selected uniformly and redrawn using a heat-bath update^15^,

$$P\left( s_{i}=+1 \right)=\frac{1}{1+exp\left( -2\beta F_{i} \right)}.$$

Exchange and turnover events are randomly interleaved within each branch^13^. The turnover rate γ is identical in SOFT and STIFF. We use this term only as a generic background epigenetic turnover process; it is not a kinetic model of DNMT activity. A γ = 0 sensitivity test removes this process entirely.

#### Simulation design, control correction, and observables

The model is read at t = 1 MCS. We treat this as a finite-time response of an already established SOFT methylation landscape rather than an equilibrium state. For every stochastic initial realization, we run three branches from the same starting configuration: SOFT (λ = 2), an independent SOFT control branch (λ = 2), and STIFF (λ = 5). J, η_cross, h_H, h_A, μ, γ, β, lattice size, and readout time are identical across the three branches. Therefore, λ is the only condition-specific model parameter changed between SOFT and STIFF.

The independent SOFT control quantifies stochastic drift over the same finite observation window. For each genomic bin, we define the branch-wise methylation changes relative to the matched SOFT branch as

$$\Delta_{\mathrm{STIFF}}\left( i \right)=m_{\mathrm{STIFF}}\left( i \right)-m_{\mathrm{SOFT}}\left( i \right), \Delta_{\mathrm{CTRL}}\left( i \right)=m_{\mathrm{CTRL}}\left( i \right)-m_{\mathrm{SOFT}}\left( i \right).$$

$$\Delta m_{X,corr}=\left\langle\Delta_{\mathrm{STIFF}} \right\rangle_{X}-\left\langle\Delta_{\mathrm{CTRL}} \right\rangle_{X}, X\in\{H,A\}.$$

The control-corrected global shift is defined analogously. To quantify local redistribution independent of sign, we use the mean absolute change and subtract the corresponding control value,

$$R_{\mathrm{corr}}=\left\langle\left| \Delta_{\mathrm{STIFF}} \right| \right\rangle-\left\langle\left| \Delta_{\mathrm{CTRL}} \right| \right\rangle.$$

**Boundary analysis-** Strong SOFT methylation-state boundaries are called from the ensemble SOFT profile. The profile is first classified into low, intermediate, and high states using the thresholds above and adjacent bins of the same state are merged into runs. A boundary is retained when both flanking runs are at least 20 kb long and their mean methylation levels differ by at least 0.15. These are methylation-state boundaries defined within the model and should not be interpreted as CTCF or TAD boundaries. For each bin, we compute the genomic distance to the nearest strong SOFT boundary. Boundary-proximal bins are defined as ≤50 kb from a boundary and domain-interior bins as ≥250 kb away. Boundary-lability excess is then calculated as

$$B=\left[ \left\langle\left| \Delta_{\mathrm{STIFF}} \right| \right\rangle_{\mathrm{near}}-\left\langle\left| \Delta_{\mathrm{STIFF}} \right| \right\rangle_{\mathrm{far}} \right]-\left[ \left\langle\left| \Delta_{\mathrm{CTRL}} \right| \right\rangle_{\mathrm{near}}-\left\langle\left| \Delta_{\mathrm{CTRL}} \right| \right\rangle_{\mathrm{far}} \right].$$

For distance-resolved profiles, bins are grouped at 0–10, 10–25, 25–50, 50–100, 100–150, 150–200, 200–250, and ≥250 kb from the nearest boundary.

**SOFT preservation and robustness analyses-** Before examining the STIFF branch, we independently verified that the primary parameter set preserves the SOFT architecture through 1 MCS. The predeclared criteria were: absolute error ≤0.01 for the H-like mean, A-like mean, and global mean methylation; absolute error ≤0.05 for the high- and intermediate-state fractions; relative error ≤25% for the mean high- and intermediate-state run lengths; and absolute error ≤10 kb for the corresponding medians. The final verification used 500 stochastic realizations and a fresh master seed that was not used during parameter selection. All criteria passed at 0.25, 0.5, and 1 MCS (Supplementary Fig. S9).

We next evaluated the frozen model across 20 independent master seeds, with 250 stochastic realizations per branch and per seed (Supplementary Fig. S10). To test the assumption of preferential same-material cooperativity, η_cross was varied over 0, 0.10, 0.25, and 0.50 using 10 master seeds and 200 stochastic realizations per branch (Supplementary Fig. S11). A confinement result was interpreted as robust only when the corresponding SOFT baseline passed the predeclared preservation criteria. We also generated 10 independent empirical-like SOFT landscapes. Each landscape matched the same state-fraction and run-length constraints but had a different spatial arrangement; methylation values within each state were resampled from the corresponding state-class values in the frozen reference track. Each landscape was evaluated with five independent master seeds and 200 stochastic realizations per branch, without landscape-specific recalibration (Supplementary Fig. S12). Finally, we varied J over 0.6, 0.8, and 1.0 and γ over 0, 0.001, 0.003, and 0.01, using 10 master seeds and 200 stochastic realizations per branch for each value (Supplementary Figs. S13 and S14).

For seed-level robustness analyses, individual master seeds are the independent replicate unit. For landscape generalization, model effects are first averaged over the five master seeds within each landscape and the 10 landscape means are used as the independent replicate unit. Summary error bars report 95% confidence intervals across these replicate units. Genomic bins are not treated as independent replicates for model inference.

**Image Analysis of H3K9me3 expression**

Line intensity profiles of H3K9me3 immunofluorescence were extracted from individual cell nuclei (n = 5 per condition) using the plot profile function in Fiji/ImageJ (NIH). For each biological replicate, a line scan was drawn across the nucleus to record raw grey values as a function of distance (micrometers). The spatial heterogeneity of H3K9me3 within each nucleus was subsequently quantified by calculating the Coefficient of Variation (CV) from the raw intensity values to capture the dispersion and relative fluctuation of grey values along the scan line, calculated as CV = (sigma / mu) x 100, where sigma is the sample standard deviation computed with Bessel's correction, and mu is the arithmetic mean grey value of the profile. Standardized CV metrics were compared between experimental groups using Welch's independent two-sample t-test without assuming equal variances.

**Image Analysis and Statistical Correlation of Collagen and AcH3K27 expression**

Mean Fluorescence Intensities (MFI) for extracellular matrix collagen and acetyl-histone H3 Lysine 27 (AcH3K27) were quantified from matched bivariate imaging datasets (n = 102 paired measurements; 51 per biological group). Normality testing using the Shapiro-Wilk method indicated that AcH3K27 expression significantly deviated from a normal distribution (P < 0.05), necessitating the use of non-parametric statistical frameworks. The monotonic relationship between pooled collagen and AcH3K27 levels across Healthy and Periodontal Disease cohorts was evaluated using a two-tailed Spearman rank-order correlation, with statistical significance defined at alpha = 0.05. Bivariate observations were visualized as a scatter plot color. A global linear regression trajectory computed via ordinary least squares was overlaid alongside a shaded 95% confidence interval, with the calculated Spearman's rank correlation coefficient (rho) and exact P-values reported directly on the figure.

**Nuclear Envelope Gene Set Curation**

A nuclear envelope gene set was assembled by querying the Gene Ontology database through the Bioconductor library (v3.19) using the following terms: nuclear envelope (GO:0005635), nuclear lamina (GO:0005652), nuclear pore complex (GO:0005643), cellular protein localisation to nucleus (GO:0034613), nuclear envelope organisation (GO:0006998), nuclear envelope targeting (GO:0070182), nuclear retention (GO:0051645), and mitotic nuclear membrane disassembly (GO:0007077). Only experimentally or biologically inferred evidence codes were accepted; electronic annotations were excluded. This yielded 217 genes, which were supplemented with a manually curated panel of 75 structural elements validated by primary literature, including A- and B-type lamins, LINC complex components, and core nuclear pore complex subunits.

Differentially expressed genes from bulk RNA sequencing of gingival fibroblasts in stiff (G' 2.0 kPa) versus soft (G' 0.75 kPa) collagen-alginate hydrogels were filtered against this gene set. Genes meeting a Benjamini-Hochberg adjusted P-value below 0.05 by DESeq2 analysis yielded 41 candidates for downstream enrichment testing.

Gene ontology biological process enrichment was performed using the clusterProfiler package (v4.x) in R with the enrichGO function, using Ensembl identifiers mapped against the org.Hs.eg.db reference database. Statistical over-representation was assessed by one-sided hypergeometric test with Benjamini-Hochberg correction, retaining terms at q < 0.05. Results were visualised as a bubble plot in ggplot2, with the x-axis showing gene ratio and the y-axis ordering terms by statistical significance. Bubble size scaled with gene count per term and colour mapped to -log10(adjusted P-value).

**Collagen Methylation Analysis**

To evaluate the epigenetic remodeling of the collagen gene family under substrate stiffness, a multi-tiered Sankey flux pathway was modeled using a whole-genome bisulfite sequencing (WGBS) dataset comparing human gingival fibroblasts cultured in stiff versus soft hydrogels. A curated cohort of 40 collagen-encoding genes (COL1A1 to COL27A1) was cross-referenced against a master differentially methylated region (DMR) table containing 13,490 functional annotation records. Bipartite directional linkages were defined to connect individual gene nodes to their dominant methylation polarity (hypermethylated or hypomethylated under stiff conditions) and subsequently routed into specific genomic feature contexts: Transcription Start Site (TSS), 5'UTR, Exon, Intron, and 3'UTR. Flux link widths were scaled proportionally to the raw frequency of overlapping DMR feature annotations per gene-direction-region combination. Genes featuring at least one statistically significant DMR within a promoter-proximal region-defined by the promoter assignment flag in the annotation pipeline-were marked with a star symbol (*) to highlight direct transcriptional regulatory networks.

**Transcription Factor Binding Motif Enrichment in Differentially Methylated Regions Analysis**

Differentially methylated regions (DMRs) in the CG context were identified from whole-genome bisulfite sequencing (WGBS) data comparing human gingival fibroblasts cultured in stiff versus soft hydrogels. DMRs were partitioned by directional methylation polarity into hypermethylated or hypomethylated cohorts, yielding 1,291 unique hypermethylated and 17,373 unique hypomethylated non-overlapping intervals across the genome. Transcription factor (TF) binding motif enrichment within these coordinates was determined via HOMER (v4.11) utilizing the findMotifsGenome.pl command against the hg38 reference assembly, with exact interval sizes enforced and genomic repeats masked. Statistical significance was calculated using a binomial test comparing target DMR frequencies against genome-wide background sequences, with raw significance values converted to Benjamini-Hochberg false discovery rate (FDR) adjusted q-values. Eight TF binding motifs demonstrated significant enrichment (q < 0.05) exclusively within the hypomethylated regions (CTCF, c-Myc, BORIS, NRF1, E2F1, E2F4, HINFP, and bHLHE41), whereas no individual motif reached statistical significance within the hypermethylated regions (data not shown). For non-canonical NF-κB pathway, subunits were identified among the scanned motifs by Rel Homology Domain (RHD) annotation: NFkB2-p52 and the NF-κB p50/p52 heterodimer. The non-canonical NF-κB pathway predominantly targets activation of the p52/RelB NF-κB complex, with pathway activation depending on the inducible processing of p100. Pathway membership was defined as NIK/MAP3K14-dependent (non-canonical) versus IKK-dependent (canonical), consistent with activation of non-canonical NF-κB involving signal-induced disruption of the cIAP E3 complex

**Chromatin Compartment Mapping and Proportional Genomic Feature Distribution of DMRs**

To determine the structural distribution of mechanical stiffness-responsive epigenetic alterations, differentially methylated regions (DMRs) pre-called pairwise from whole-genome bisulfite sequencing (WGBS) data of human gingival fibroblasts (stiff versus soft hydrogels) were mapped to the hg19 genome assembly. Overlapping genomic coordinates were deduplicated to retain 1,294 unique hypermethylated DMRs and 17,390 unique hypomethylated DMRs. Concurrently, the autosomal genome was partitioned into non-overlapping bins at 3.276 Mb resolution and assigned chromatin compartments. Bins were classified by a majority base-pair rule (greater than 50% span) into heterochromatin, euchromatin, and intermediate states (263 bins). Individual DMR midpoints were mapped to these intervals achieving assignment rates of 95.7% for hyper-DMRs and 98.5% for hypo-DMRs. Analysis was restricted to heterochromatin-compartment bins, retaining 646 hyper-DMRs and 7,054 hypo-DMRs after filtering missing annotations. Ten raw genomic reference categories were consolidated into six functional feature groups: (1) Repeat, (2) Intron, (3) CpG island/shore, (4) Promoter/TSS, (5) Exon/5'UTR, and (6) 3'UTR/TES. The proportional feature composition of each methylation class was modeled into a 10 x 10 mosaic waffle grid where each tile represents 1% of the annotated dataset. Integer rounding residuals were adjusted at the most abundant category (Repeat), while elements falling below the 0.5% detection threshold (3'UTR/TES) received zero tiles. Quantitative distribution profiling revealed that repeat elements comprised 61% of hyper-DMRs (n = 393) compared to 46% of hypo-DMRs (n = 3,278), whereas introns and CpG islands/shores constituted 24% and 13% of hypo-DMRs versus 18% and 9% of hyper-DMRs, respectively.

**Genome-wide Visualization of Compartment-Stratified CpG Methylation Change**

Chromatin Compartment-Level Whole-Genome Bisulfite Sequencing Analysis To evaluate the macro-scale structural distribution of mechanosensitive DNA methylation shifts, CpG context methylation profiles from whole-genome bisulfite sequencing (WGBS) data of human gingival fibroblasts (stiff versus soft hydrogels) were quantified across 854 non-overlapping autosomal bins at a 3.276 Mb resolution. The methylation differential (Delta-beta) was defined as CG_Stiff - CG_Soft, where positive values denote a net methylation gain under stiff substrate conditions. Bins were mapped to chromatin compartments by intersecting coordinates with the Roadmap Epigenomics 15-state ChromHMM model. Chromatin states were collapsed into heterochromatin, euchromatin, and intermediate classifications, with compartment assignment governed by a majority base-pair rule requiring greater than 50% coverage of the genomic span. Bins failing to meet this threshold were categorized as unassigned. This stratification isolated 542 heterochromatin, 49 euchromatin, 244 unassigned, and 19 intermediate bins. To isolate high-confidence structural environments, downstream visualization was restricted to validated heterochromatin and euchromatin states (n = 591).

For genome-wide spatial tracking, a horizontal Manhattan layout was constructed along a continuous genomic axis by concatenating chromosomes 1-22 with a 10 Mb inter-chromosome spacer, mapping each bin relative to its structural midpoint. Heterochromatin entries were represented as filled circles and euchromatin entries as filled triangles, referenced against a horizontal baseline at y = 0. Alternatively, a vertical layout was implemented by arranging individual chromosomes as stacked horizontal strips, where the horizontal axis encoded the absolute Delta-beta magnitude and within-strip bin positions were jittered by +/- 0.15 to mitigate overplotting. Quantitative analysis revealed a 2.11 percentage point (pp) compartment separation, characterized by a mean methylation gain in heterochromatin (+0.959 pp) and a mean reduction in euchromatin (-1.154 pp). This coordinated behavior was further underscored by strong compartment-level directionality, with 85.4% of heterochromatin bins exhibiting positive trajectories and 93.9% of euchromatin bins exhibiting negative trajectories. The widest variation amplitudes were concentrated within heterochromatic domains, spanning a range from -6.66 to +7.78 pp, compared to a tighter euchromatin range of -3.97 to +0.41 pp.

**Nuclear Lamin B1 expression Analysis**

Confocal z-projections of Lamin B1 immunofluorescence (Alexa Fluor 594; Imaris Viewer 11.0.0) were exported as 8-bit RGB TIFF files. The red channel was extracted and a diffuse background estimate was subtracted by computing a large-scale Gaussian-smoothed image (σ = 50 px) and subtracting it from the raw signal, clipping negative values to zero. The result was smoothed with a Gaussian kernel (σ = 2 px) to reduce photon shot noise while preserving fold-scale features. Nucleus segmentation used three-class Otsu thresholding applied to a separately smoothed copy of the raw image (σ = 4 px), yielding two intensity boundaries. The lower threshold (t_low) defined the full Lamin B1 signal mask (nucleus shape); the upper threshold (t_high, ~2% of image area) isolated the densest ring pixels. Small disconnected objects were removed (< 500 px²) and binary closing was applied (disk radius 5 px for the shape mask, 3 px for the ring mask).

**Curvature-coloured contour maps-** The nucleus boundary contour was extracted from the t_low shape mask using the marching squares algorithm (iso-level 0.5; skimage.measure.find_contours). Local curvature κ was estimated at each image pixel as the negative Laplacian-of-Gaussian of the background-subtracted image (σ = 3 px): κ(x,y) = −∇²[G_σ * I](https://claude.ai/chat/x,y), where positive values correspond to convex intensity peaks (fold apices) and negative values to concave regions (invaginations). Curvature was sampled at contour coordinates and smoothed along the arc (Gaussian, σ = 20 contour points) for visualization. The contour was rendered as a colour-mapped line (RdBu_r, diverging colormap centred at zero) using matplotlib LineCollection, with the colour range clipped to the 2nd–98th percentile of the per-nucleus curvature distribution. An ellipse residual contour was additionally computed by fitting a reference ellipse to the contour (algebraic least-squares; skimage.measure.EllipseModel) and computing the signed radial deviation of each contour point from the ellipse boundary (positive = protrusion, negative = invagination; PuOr_r colormap). Colourmaps are normalised independently per nucleus; the scalar metrics (mean |κ|, excess perimeter ratio) are used for quantitative cross-nucleus comparison.

**Structure tensor coherence-** The structure tensor J = G_σ * (∇I ⊗ ∇I) was computed on the background-subtracted smoothed image with σ = 3 px (skimage.feature.structure_tensor, order = 'rc'). The coherence index C at each pixel was derived analytically from the tensor components (A_xx, A_xy, A_yy) without explicit eigendecomposition: C(x,y) = √[(A_xx − A_yy)² + 4A_xy²] / (A_xx + A_yy + ε), where ε = 10⁻⁹ prevents division by zero. C ranges from 0 (isotropic gradient field; flat or noisy lamina) to 1 (perfectly linear intensity edge; sharp fold). The dominant local fold orientation was computed as θ = 0.5 × arctan2(2A_xy, A_xx − A_yy), giving the fold axis direction in [0°, 180°]. Because orientations are axial, circular statistics were performed on doubled angles. The coherence-weighted mean resultant length R̄ = √[(Σ w_i cos 2θ_i)² + (Σ w_i sin 2θ_i)²], with weights w_i = C_i / ΣC_j, quantifies directional order: R̄ = 0 indicates randomly oriented folds; R̄ = 1 indicates complete alignment. Circular variance was defined as 1 − R̄.

**Angular ring intensity profile-** The intensity-weighted centroid of each nucleus was computed from the shape mask. A dynamic ring zone was identified by computing a 100-bin radial intensity profile from the centroid and locating the peak radius r_peak (the radius of maximum mean intensity). The analysis zone was defined as r_peak ± 15%. The circumferential profile was sampled at 1080 uniformly spaced angles (0.33° resolution); at each angle, the maximum intensity along a radial transect of 60 points spanning the ring zone was recorded. This max-projection approach captures inward folds regardless of their exact radial position within the zone. The profile was smoothed with a Gaussian filter (σ = 2.0 sample points) and peaks were detected using an adaptive prominence threshold: prom_min = max(μ_P × CV_ring × 3, 0.08 × (P_max − P_min)), where CV_ring = σ(P)/μ(P) is the profile coefficient of variation. This suppresses noise peaks in smooth nuclei (low CV) while remaining sensitive to true wrinkle peaks in folded nuclei (high CV). Wrinkle wavelength λ was estimated as the mean inter-peak arc length. The dimensionless bending stiffness-to-tension ratio B/σ was computed following Cerda and Mahadevan (2003) as B/σ = (λ/2πr_peak)², providing a mechanical proxy for the lamina's wrinkling state.

All analyses were implemented in Python 3.12 using NumPy, SciPy, scikit-image, Pillow, and Matplotlib. This analysis pipeline was developed with the assistance of Claude (Anthropic, claude.ai), an AI assistant, which aided in code structure, mathematical implementation, and figure generation. All biological interpretations, parameter choices, and validation were performed by the authors.

**CUT&RUN Sequencing**

CTCF-bound DNA from GFs was extracted using the CUT&RUN Assay Kit (Cell Signaling Technology), monoclonal rabbit anti-CTCF ([confirm clone, e.g. D31H2]), and DNA Purification Buffers and Spin Columns (Cell Signaling Technology) after cell retrieval from the matrix. Normal rabbit IgG was used as a negative control, and S. cerevisiae spike-in DNA (Cell Signaling Technology) was added to each reaction for normalization. Libraries for next-generation sequencing were prepared using 1 ng of CUT&RUN DNA and the NEB Ultra II DNA library preparation kit (New England Biolabs, Ipswich, MA) according to the manufacturer’s instructions, with 12 cycles of final amplification. Library size was assessed using the 4200 TapeStation and the High-Sensitivity DNA assay (Agilent, Santa Clara, CA). Concentration was determined using the Qubit Fluorometer 2.0 (Thermo Fisher, Waltham, MA). Libraries were pooled and sequenced with a 61 × 61 bp run length on the NextSeq 2000 (Illumina, San Diego, CA). A minimum of 26 million read pairs was acquired for each sample. Raw sequencing reads were processed using the nf-core/cutandrun pipeline (v3.2.2) in Nextflow. Reads were trimmed with Trim Galore and aligned to the human (GRCh38) and yeast (R64-1-1) genomes using Bowtie2. Peaks were called with MACS2 in paired-end mode against the IgG control (q < 0.01), after removal of blacklisted regions. Analyses were performed in R (v4.6.0). Peak summits (±200 bp) were merged across samples, and sites called in at least two of six samples were retained (63,844 sites). Shared sites were called in at least one replicate of each condition (44,780 sites). Read pairs per site were counted with Rsubread and normalized to yeast spike-in read pairs. Libraries were prepared on two occasions, and spike-in size factors were calculated relative to the libraries prepared together. For each replicate, mean normalized signal across sites in a group was expressed relative to the soft mean. Heatmaps were generated with EnrichedHeatmap and ComplexHeatmap. Genomic regions were annotated with ChIPseeker (promoter, ±3 kb of the TSS). CTCF motifs were identified using the JASPAR MA0139.2 matrix (Biostrings, P < 10⁻⁴). For 24,266 motif-containing shared sites, fragment midpoints within ±500 bp of the motif were plotted against fragment length (Rsamtools), oriented by motif strand and normalized to spike-in. Fragments ≤120 bp were assigned to the CTCF footprint and 150–260 bp fragments to flanking nucleosomes. CG-context DMRs from WGBS were intersected with CTCF sites. Background windows were sampled from the same chromosome and matched to each DMR for width, GC content, and CpG density. Expected overlap and empirical P-values were derived from 10,000 matched draws. Methylation change and CTCF change were compared by Spearman correlation. LADs (HFFc6 lamin B1 DamID; 4DNFIN63IWZ3, 4DNFIHHIVIME) and TAD boundaries (HFFc6 Micro-C; 4DNFIC8438YF) were obtained from the 4D Nucleome Data Portal. LADs present in both replicates and ≥10 kb were retained. CTCF site density around LAD borders and TAD boundaries (±100 kb, 5 kb bins) was expressed relative to 500,000 random genomic positions. Replicate-level CTCF occupancy was compared between soft and stiff conditions by two-tailed unpaired t-test (n = 3)

**Statistical tests, software, and use of artificial intelligence**

Data are presented as mean ± standard deviation from at least three biologically independent donors or experiments. All datasets were tested for normality prior to parametric analysis. Differences between two groups were assessed by two-tailed unpaired or paired t-test, with Welch's correction applied for unequal variances where appropriate. For comparisons involving more than two groups, one-way or two-way ANOVA was performed followed by Tukey's, Dunnett's, or Šídák's multiple comparisons test, the Benjamini-Hochberg FDR method, or uncorrected Fisher's LSD as indicated for each experiment. A P-value below 0.05 was considered statistically significant. Statistical analyses and graph preparation were performed in Prism (Dotmatics). Image analysis was performed in Fiji (NIH) and Imaris (Oxford Instruments). Pathway and gene set enrichment analyses were performed using Reactome and NIH DAVID. Gene ontology enrichment analysis of differentially expressed genes was performed using clusterProfiler (v4.x) in R with the enrichGO() function against the org.Hs.eg.db reference database, as described above. Whole-genome bisulfite sequencing data were processed using Bismark for alignment and DSS for differentially methylated region calling. Visualization was performed using ggplot2 and circlize in R. Bioinformatics pipelines were implemented through R and The Dry Lab platform (thedrylab.com). The schematic in Fig. 1A (top right) was generated using Biorendor and Gemini Pro (Google DeepMind). Artificial intelligence tools were used for English language editing during manuscript preparation.

**Supplementary Results**

**Computational model validation and robustness**

We first asked whether the frozen model preserves the heterogeneous SOFT methylation architecture over the finite observation window used for the confinement test. Across 500 stochastic realizations, the global mean methylation, H-like and A-like means, methylation-state fractions, and domain-length statistics remained within the predeclared acceptance criteria through 1 MCS. At the final readout, the correlation with the initial SOFT profile remained 0.962, while the global mean changed by only 7.3 × 10⁻⁵ from the target value (Supplementary Fig. S9). Thus, the SOFT reference remains stable on the timescale over which the confinement response is evaluated.

We then repeated the frozen SOFT-control-STIFF comparison across 20 independent master seeds. The response was reproducible across all seeds: the control-corrected H-like response was +0.02016 (95% CI +0.01986 to +0.02046), whereas the A-like response was −0.04411 (95% CI −0.04477 to −0.04344). Local redistribution increased by 0.01352 (95% CI 0.01301 to 0.01403), and boundary-lability excess was positive in all 20 seeds, with a mean of 0.01295 (95% CI 0.01168 to 0.01422). In contrast, the control-corrected global mean shift remained small at 7.5 × 10⁻⁵ (95% CI 5.5 × 10⁻⁵ to 9.6 × 10⁻⁵) (Supplementary Fig. S10). The confinement response is therefore dominated by redistribution across the heterogeneous landscape rather than by a large change in the genome-wide mean.

The primary model assumes preferential same-material cooperativity (η_cross = 0). To determine whether the response requires a strictly zero cross-material interaction, we increased η_cross without recalibrating any other parameter. The SOFT baseline remained valid for all 10 master seeds at η_cross = 0 and 0.10, while the same confinement response was retained. At η_cross = 0.25, only 7 of 10 SOFT baselines satisfied the predeclared preservation criteria, and at η_cross = 0.50 none did. We therefore interpret the model as requiring strongly preferential, rather than necessarily perfectly exclusive, same-material cooperativity. The higher η_cross values are shown as diagnostic sensitivity tests and are not used as valid robustness evidence once the SOFT reference itself is no longer preserved (Supplementary Fig. S11).

We next tested whether the response depended on one particular arrangement of the synthetic SOFT landscape. Ten independent empirical-like landscapes were generated using the same methylation-state fractions and run-length constraints, but with different spatial organization. Pairwise correlations between the starting landscapes were centered near zero (median r = 0.011; range −0.108 to +0.105). All 50 landscape-by-seed SOFT baseline validations passed. Across the 10 landscape means, the H-like response was +0.01742 (95% CI +0.01645 to +0.01840), the A-like response was −0.03762 (95% CI −0.04081 to −0.03443), local redistribution increased by 0.01079 (95% CI 0.01007 to 0.01150), and boundary-lability excess was +0.00852 (95% CI +0.00734 to +0.00970). The control-corrected global shift remained 9.0 × 10⁻⁵ (95% CI 7.3 × 10⁻⁵ to 1.07 × 10⁻⁴) (Supplementary Fig. S12). The response therefore generalizes across spatially distinct SOFT methylation architectures without landscape-specific parameter fitting.

Finally, we tested the two principal kinetic/energetic parameters around their primary values. Varying the cooperative interaction strength over J = 0.6, 0.8, and 1.0 preserved the SOFT baseline and retained the H-like gain, A-like loss, local redistribution, and boundary-proximal lability at all three values (Supplementary Fig. S13). Varying the background turnover rate over γ = 0, 0.001, 0.003, and 0.01 produced the same qualitative response, and all four conditions preserved the SOFT baseline (Supplementary Fig. S14). In particular, the directional response persists at γ = 0. This shows that the core confinement response is generated by the contact/exchange sector of the model and does not require the generic background turnover process. Together, these analyses support a reduced physical mechanism in which a broader and denser effective contact landscape is sufficient to redistribute methylation across an already heterogeneous SOFT architecture. The model does not assign this response to specific genes, repeat families, CTCF sites, or experimentally annotated chromatin compartments.

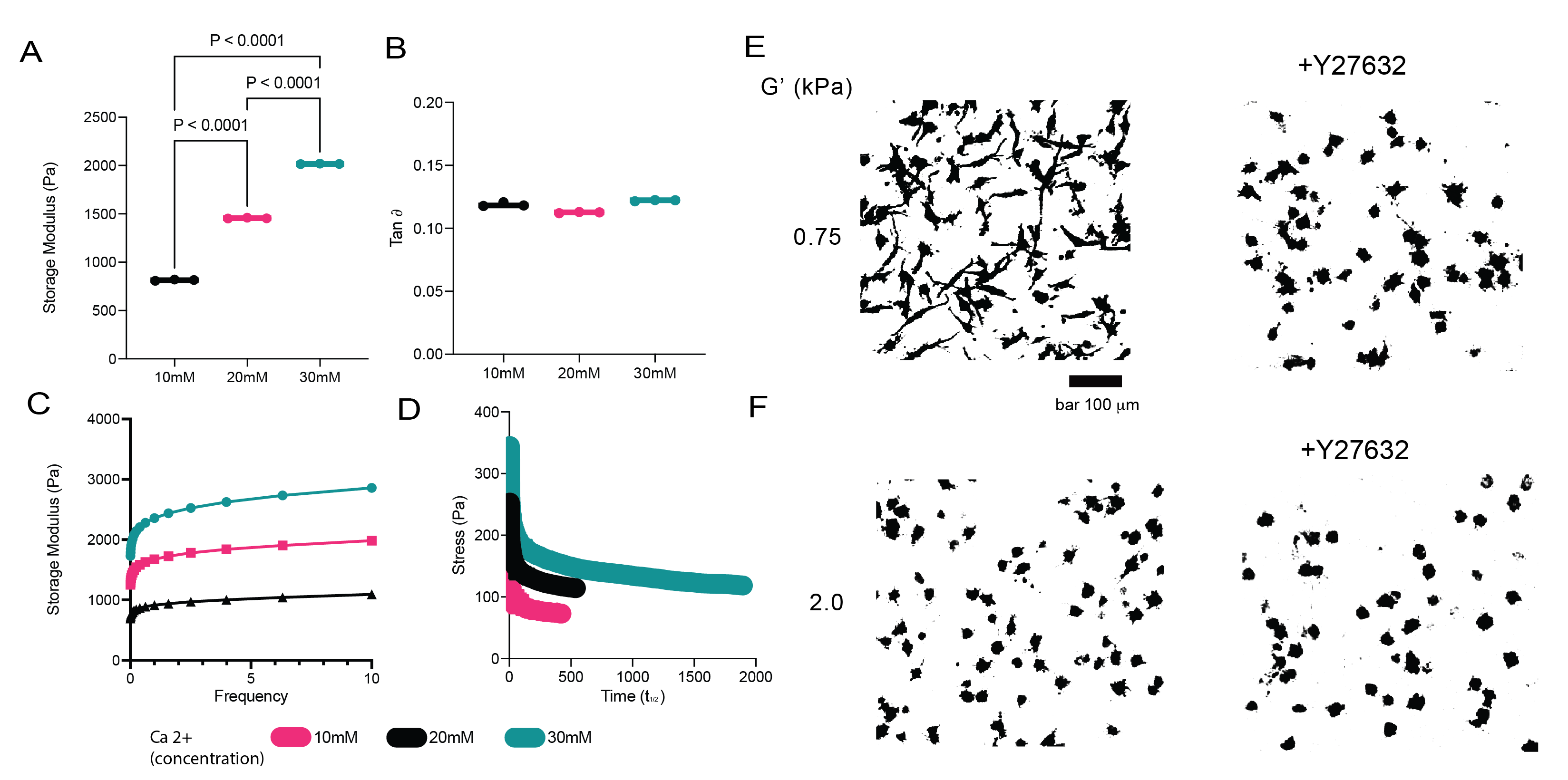

**Supplementary Figure S1.** (A) Oscillatory shear rheology- storage modulus (B) tan delta (C) frequency sweep and (D) stress relaxation behavior of gingival ECM hydrogels Data are shown as mean ± SD. P-values from statistical tests are indicated. **(E, F)** Representative binarized images of gingival fibroblasts in soft (G’ 0.75 kPa; E) and stiff (G’ 2.0 kPa; F) hydrogels, with or without Y-27632.

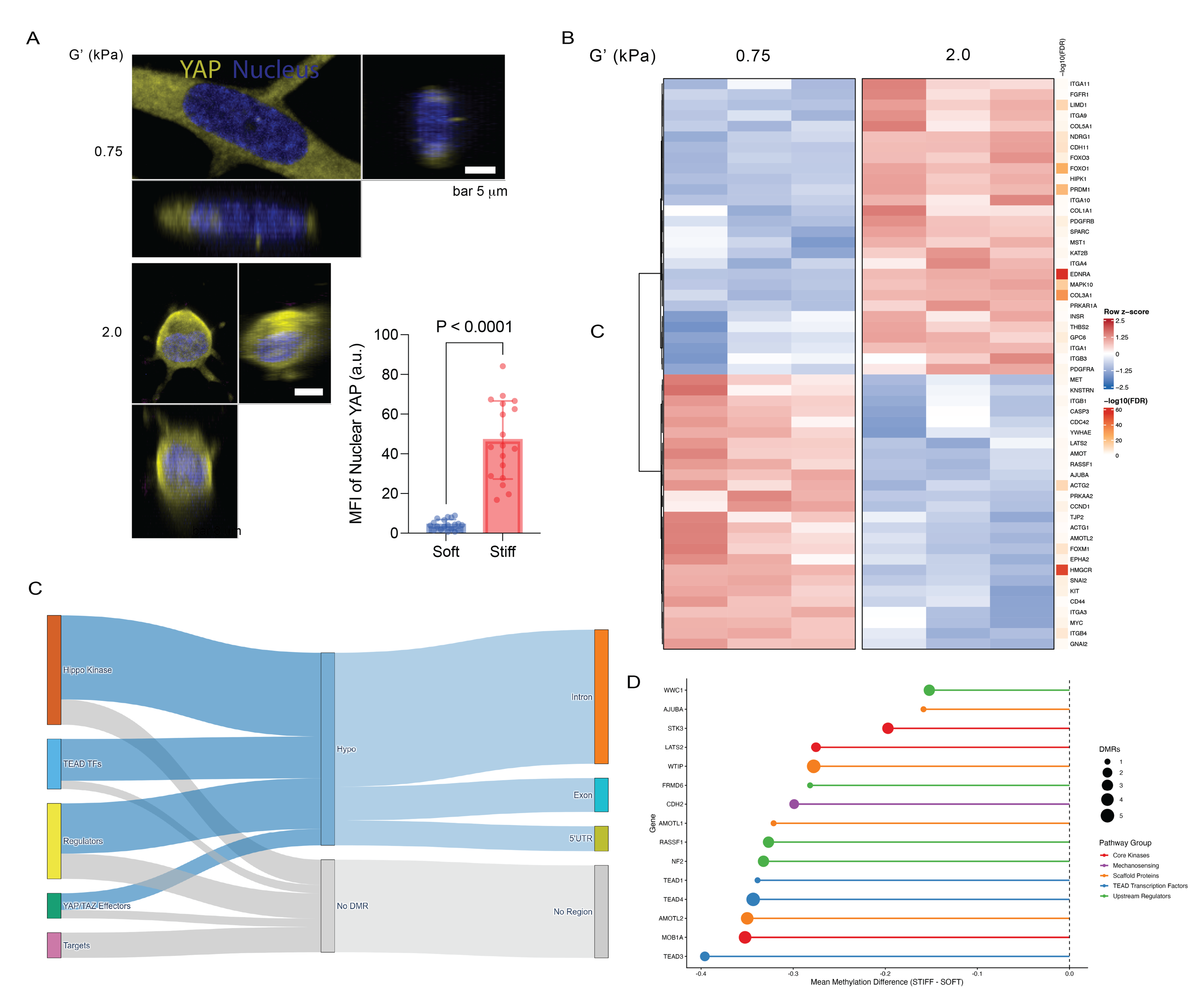
**Supplementary Figure S2. Matrix stiffness drives partial YAP nuclear translocation and hypomethylation of the YAP/Hippo pathway in gingival fibroblasts.** (A) Representative confocal immunofluorescence images of YAP (yellow) and nucleus (blue) in GFs encapsulated in soft (G' 0.75 kPa) and stiff (G' 2.0 kPa) hydrogels, with orthogonal views. Scale bars, 5 µm. Quantification of nuclear YAP mean fluorescence intensity is shown (n = 3 biologically independent donors). Data are presented as mean ± s.d. P-value was determined by two-tailed unpaired t-test. (B) Heatmap of z-scored expression of YAP/Hippo pathway genes in GFs in soft (G' 0.75 kPa) and stiff (G' 2.0 kPa) hydrogels from bulk RNA sequencing (n = 3 biologically replicates). Colour scale indicates row z-score; sidebar indicates −log10(FDR). (C) Sankey diagram of YAP/Hippo pathway differentially methylated regions in the stiff condition, grouped by functional category (Hippo kinases, TEAD transcription factors, regulators, YAP/TAZ effectors, targets), DMR direction, and genomic region. (D) Lollipop plot of the mean methylation difference (stiff minus soft) at individual YAP/Hippo pathway gene loci. Dot size indicates the number of DMRs per gene; colour indicates pathway group. Negative values indicate hypomethylation in the stiff condition.

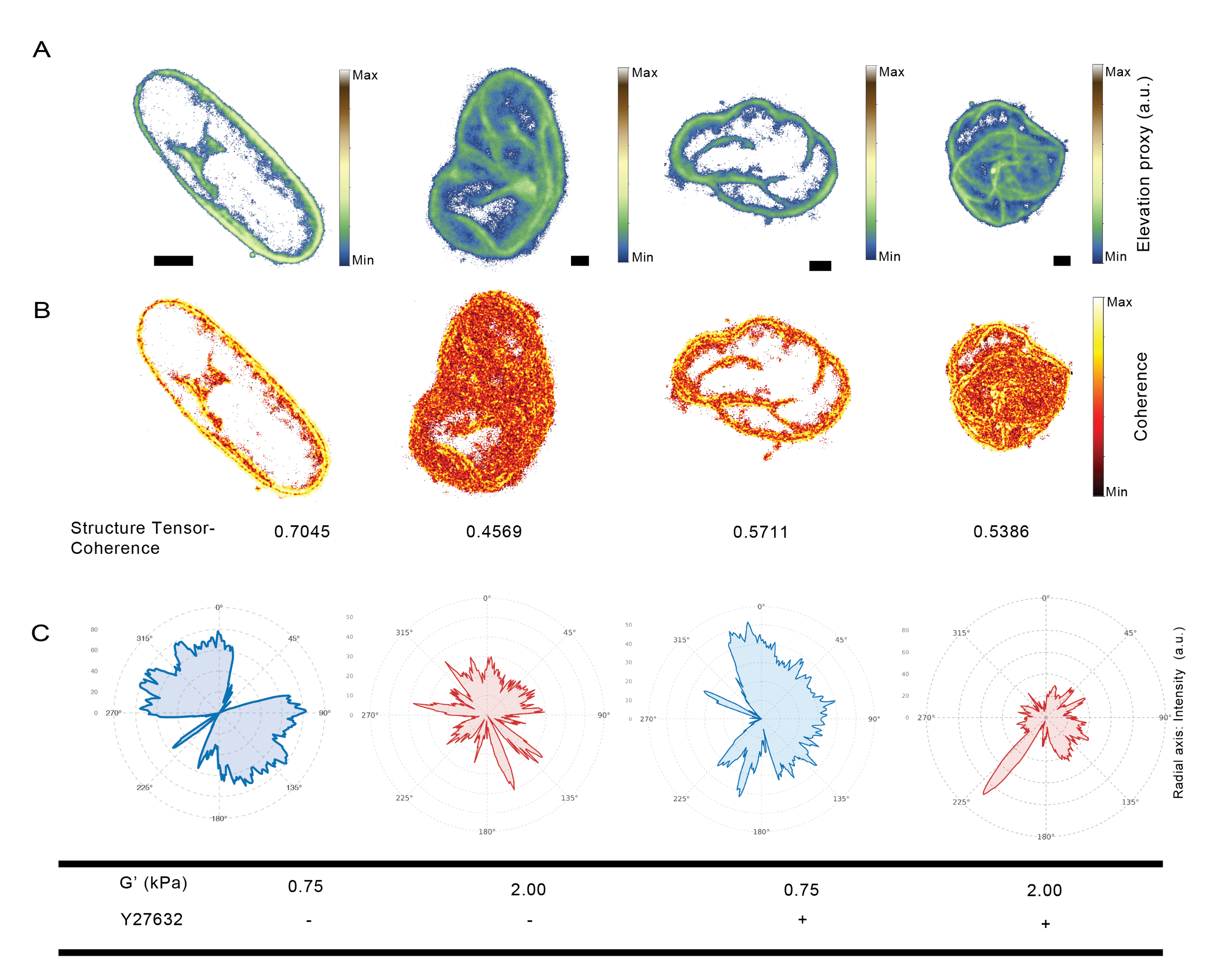

**Supplementary Figure S3. Nuclear envelope terrain mapping and structure tensor coherence analysis resolve two mechanistically distinct folding architectures across stiffness conditions and following ROCK inhibition.** (A) Lamin B1 elevation proxy maps for GFs in soft (G' 0.75 kPa) and stiff (G' 2.0 kPa) hydrogels, untreated or treated with Y-27632. Warm colours indicate regions of high Lamin B1 signal density corresponding to fold apices; cool colours indicate thin or sparse lamina regions. Scale bars as indicated. (B) Structure tensor coherence maps for the same conditions. Coherence index C = (λ₂ − λ₁)/(λ₂ + λ₁), where λ₁ and λ₂ are the minor and major eigenvalues of the local intensity gradient tensor. Score of 0 indicates isotropic gradient with no preferred fold orientation; score of 1 indicates a perfectly linear dominant fold axis. Mean coherence values are indicated below each image. (C) Polar angular Lamin B1 intensity profiles sampled at 1° increments along a circumferential ring at 70% of maximum nuclear radius for each condition. Radial axis indicates intensity in arbitrary units. G' (kPa) and Y-27632 treatment status are indicated.

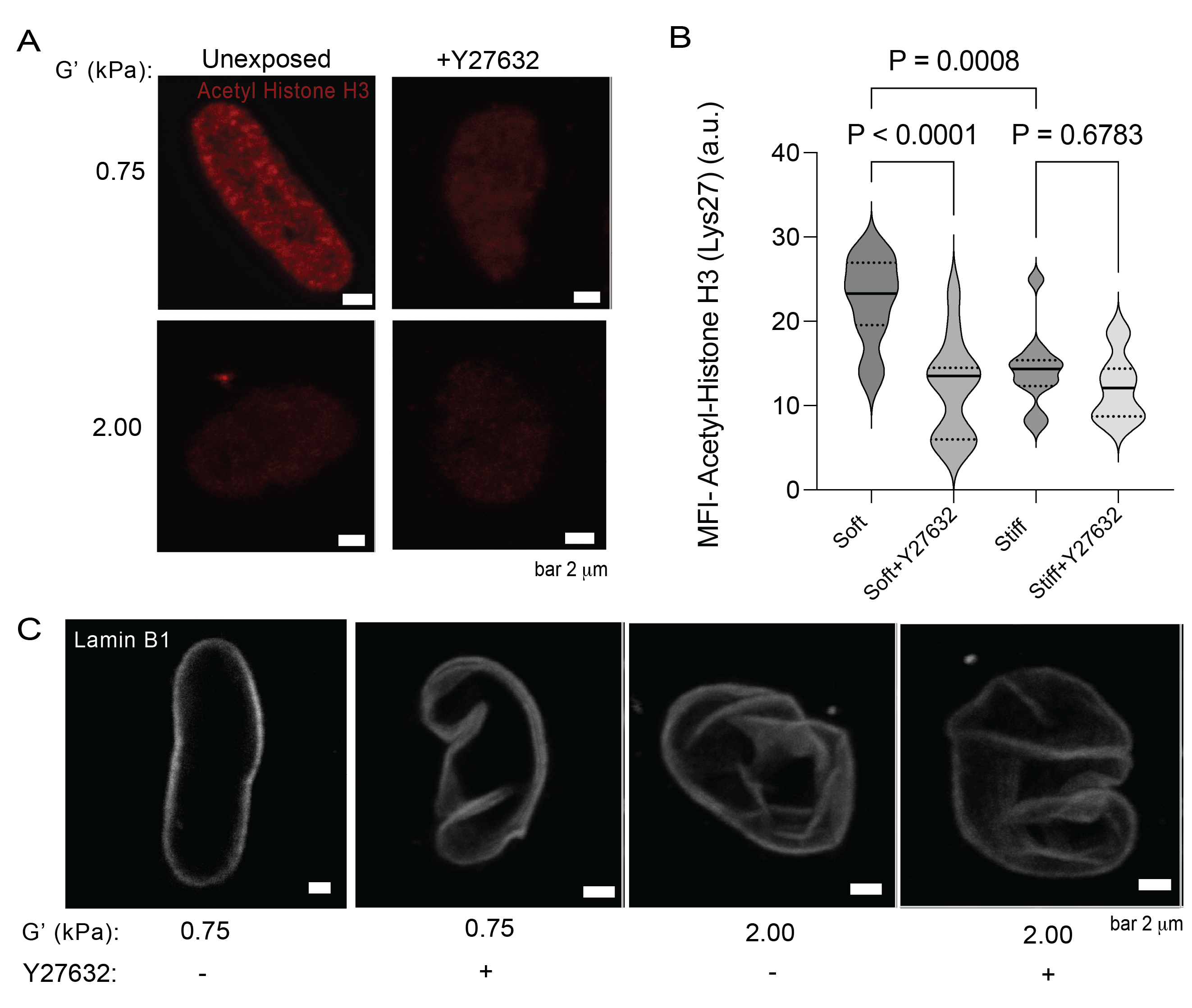

**Supplementary Figure S4. Stiffness-dependent H3K27Ac and nuclear morphology are preserved in BJ fibroblasts.** **(A)** Representative immunofluorescence images of Acetyl-Histone H3 (Lys27; red) in BJ fibroblasts cultured on soft (G’ 0.75 kPa) or stiff (G’ 2.00 kPa) hydrogels, with or without the ROCK inhibitor Y-27632. Scale bars are indicated on images. **(B)** Quantification of mean fluorescence intensity (MFI) of Acetyl-Histone H3 (Lys27) across stiffness and treatment conditions. Statistical comparisons by two-way ANOVA with Šídák correction; P-values indicated. a.u., arbitrary units. **(C)** Representative immunofluorescence images of Lamin B1 in BJ fibroblasts on soft (G’ 0.75 kPa) and stiff (G’ 2.00 kPa) hydrogels, with or without Y-27632. Smooth, elongated nuclear envelopes in soft conditions acquire folding upon Y-27632 treatment, while stiff-matrix nuclei exhibit constitutively compacted, folded morphology that is unaltered by Y-27632.

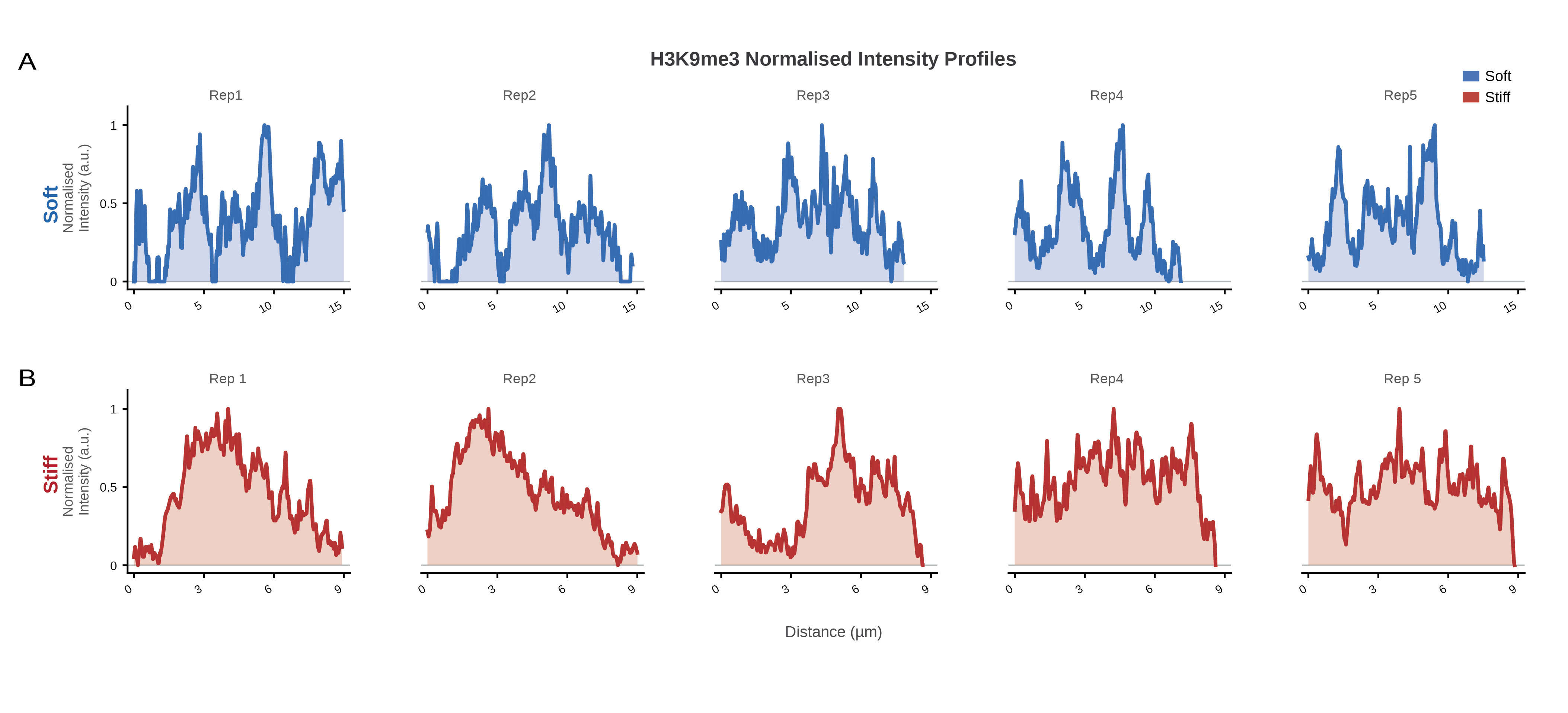

**Supplementary Figure S5. H3K9me3 normalised intensity profiles across biological replicates confirm redistribution from discrete foci to a diffuse nuclear distribution in confined fibroblasts.** (A) Normalized H3K9me3 fluorescence intensity profiles across the nuclear diameter for five biological replicates of GFs in soft (G' 0.75 kPa) hydrogels. Multi-peak, high-amplitude profiles reflect spatially clustered heterochromatin architecture. (B) Normalized H3K9me3 fluorescence intensity profiles for five biological replicates of GFs in stiff (G' 2.0 kPa) hydrogels. Broad, plateau-like profiles reflect redistribution of H3K9me3 to a spatially uniform nuclear domain. Distance in µm; normalized intensity in arbitrary units.

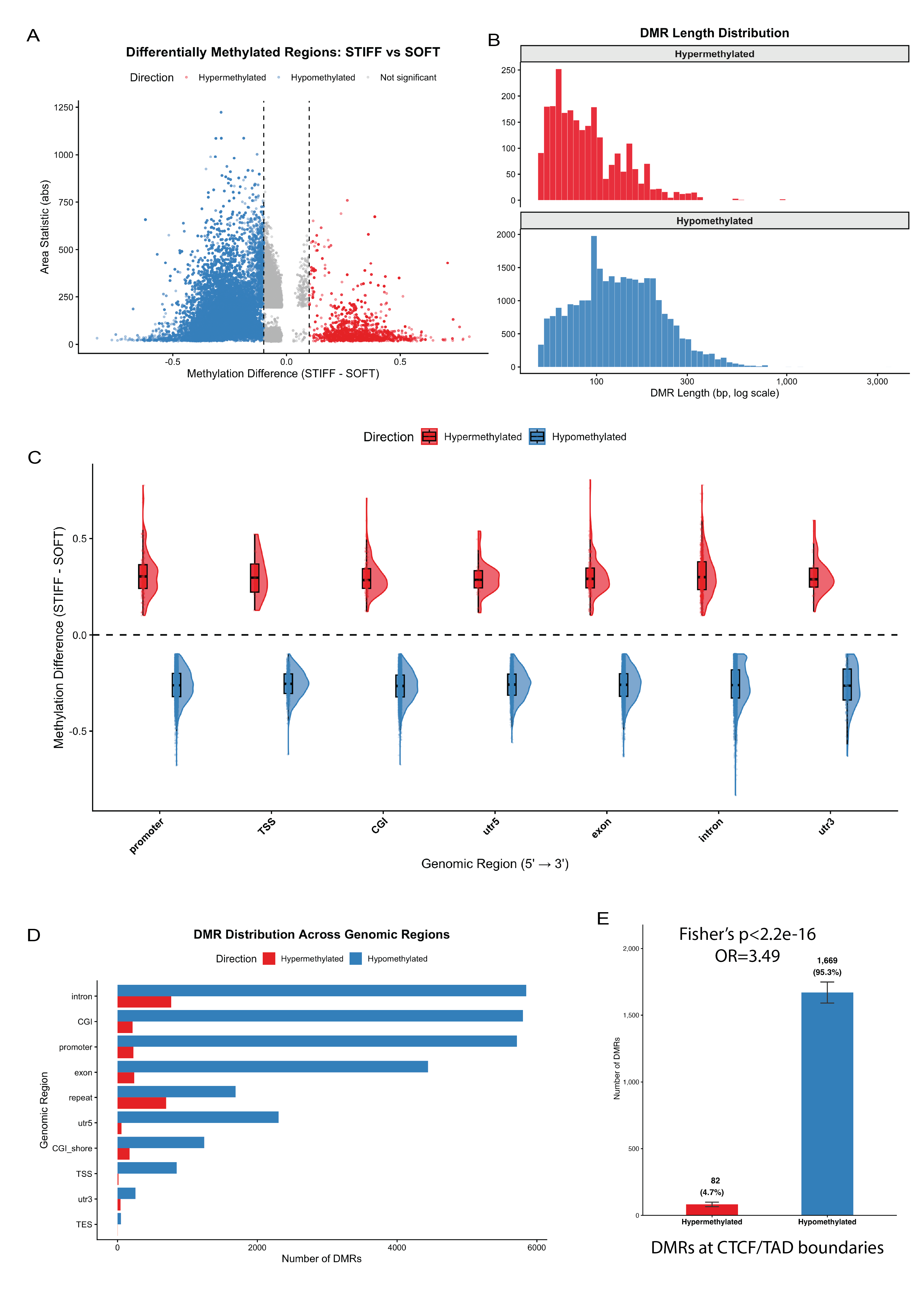

**Supplementary Figure S6. Matrix stiffness drives predominantly localised CpG hypomethylation concentrated at CTCF and TAD boundaries in gingival fibroblasts.** (A) Distribution of differentially methylated regions (DMRs) between stiff and soft conditions, plotted as absolute area statistic against methylation difference (stiff minus soft). Hypermethylated DMRs (red) and hypomethylated DMRs (blue) passing significance thresholds are shown, with non-significant regions in grey. (B) Length distribution of hypermethylated and hypomethylated DMRs on a log scale. Hypomethylated DMRs are more numerous across the size range. (C) Split violin plots of methylation difference (stiff minus soft) for hypermethylated (red) and hypomethylated (blue) DMRs across genomic regions, ordered 5' to 3' (promoter, TSS, CpG island, 5'UTR, exon, intron, 3'UTR). Box plots show median and interquartile range.

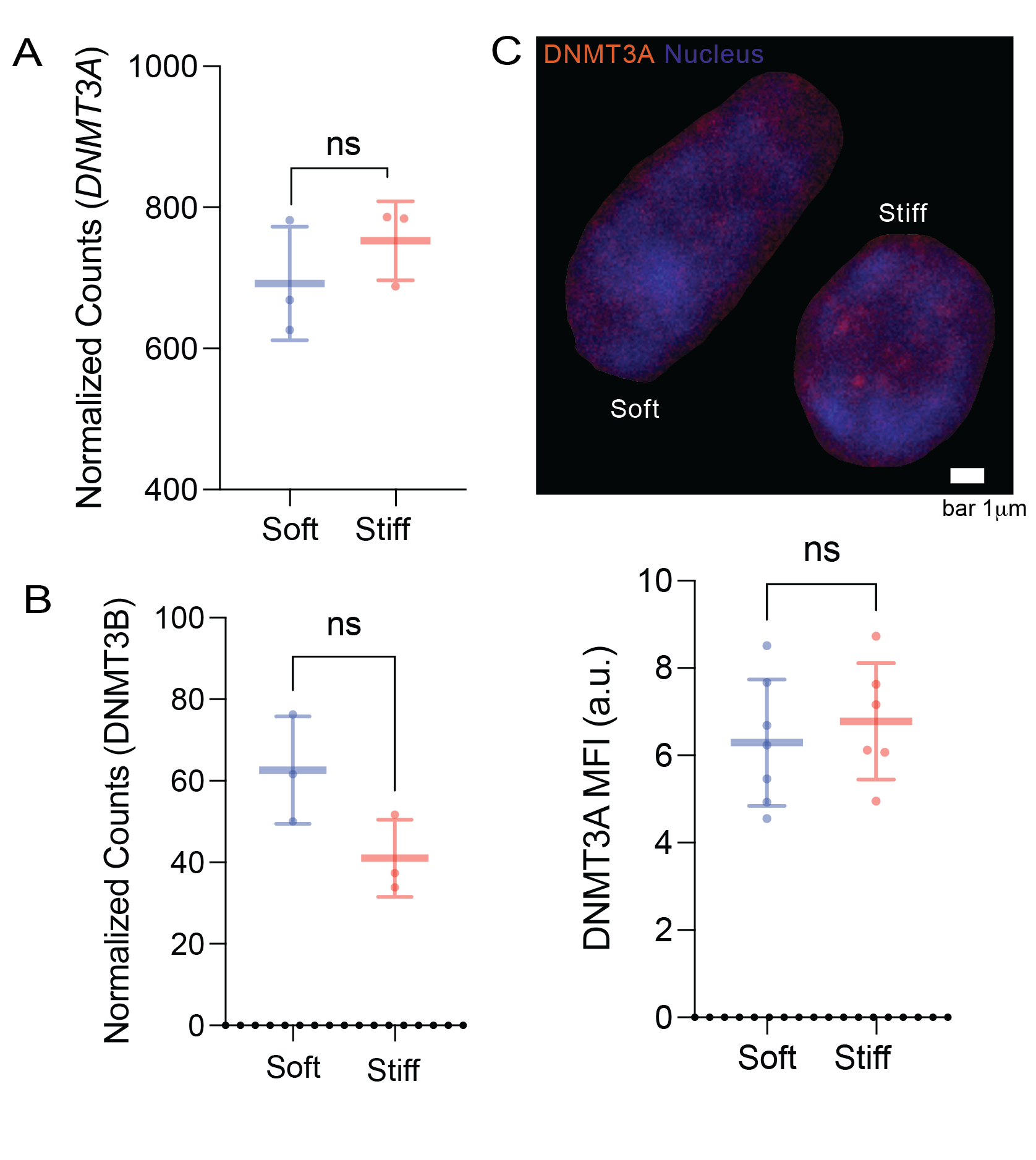

**Supplementary Figure S7. Matrix stiffness dependent expression of DNMT3A and DNMT3B (**A) Normalized RNA-seq counts for DNMT3A in gingival fibroblasts encapsulated in soft and stiff collagen-alginate hydrogels (mean ± s.d.; ns, not significant, unpaired t-test). (B) Normalized RNA-seq counts for DNMT3B under the same conditions (mean ± s.d.; ns, not significant). (C) Representative confocal images of DNMT3A immunofluorescence (orange) and DAPI nuclear stain (blue) in soft- and stiff-matrix fibroblasts. Scale bar, 1 μm. Right, quantification of DNMT3A mean fluorescence intensity (MFI) per nucleus across conditions (mean ± s.d.; ns, not significant, unpaired t-test).

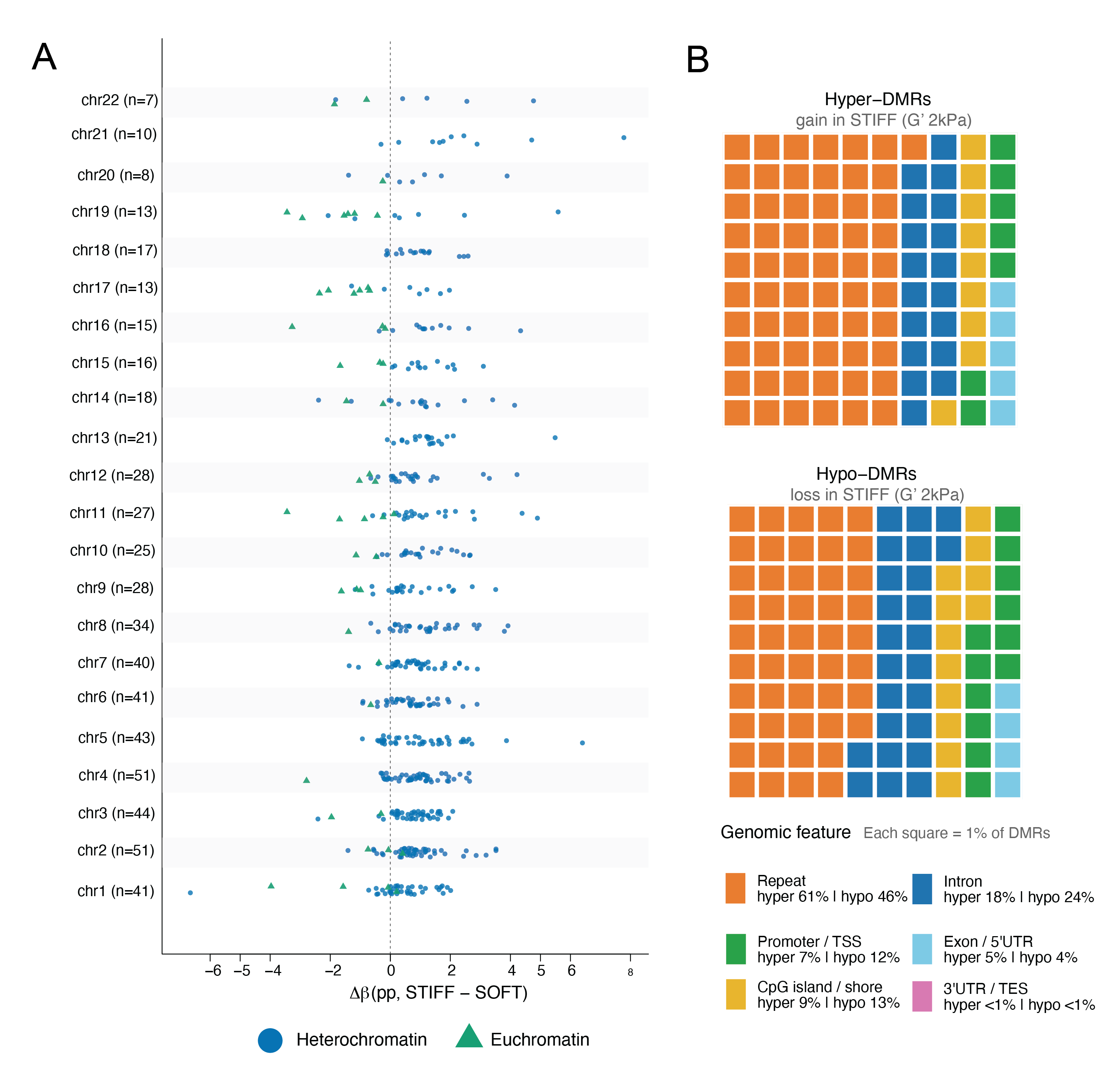

**Supplementary Figure S8. Genome-wide Δβ distribution and genomic feature composition of differentially methylated regions confirm a bifurcated methylation programme across all autosomes.** (A) Dot plot showing the distribution of methylation change (Δβ, stiff minus soft) across all autosomes for heterochromatin-associated (blue circles) and euchromatin-associated (green triangles) DMRs. Rightward shift indicates methylation gain in stiff; leftward shift indicates methylation loss. (B) Waffle chart showing the genomic feature distribution of hypermethylated DMRs (gain in stiff, G' 2 kPa) and hypomethylated DMRs (loss in stiff, G' 2 kPa). Each square represents 1% of total DMRs. Feature categories: Repeat (orange), Intron (blue), CpG island/shore (yellow), Promoter/TSS (green), Exon/5'UTR (light blue), 3'UTR/TES (pink).

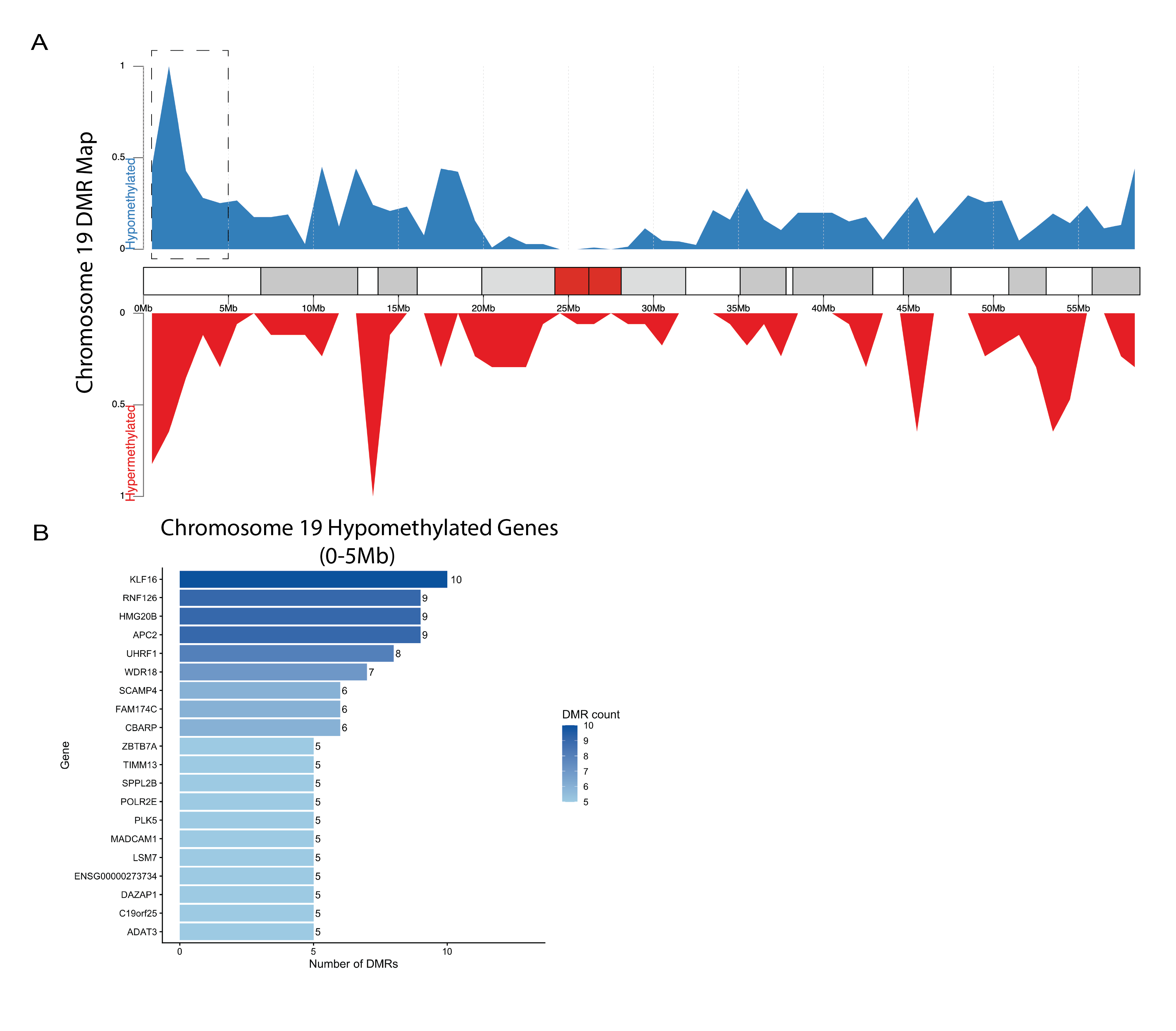

**Supplementary Figure S9. Chromosome 19 carries the highest density of differentially methylated regions, concentrated in a gene-dense hypomethylated region.** (A) Distribution of hypomethylated (blue, top) and hypermethylated (red, bottom) differentially methylated region density along chromosome 19, aligned to the chromosome ideogram. The 0-5 Mb region (dashed box) shows the highest hypomethylated DMR density. (B) Top hypomethylated genes within the chromosome 19 0-5 Mb region, ranked by the number of hypomethylated DMRs per gene. Colour scale indicates DMR count per gene.

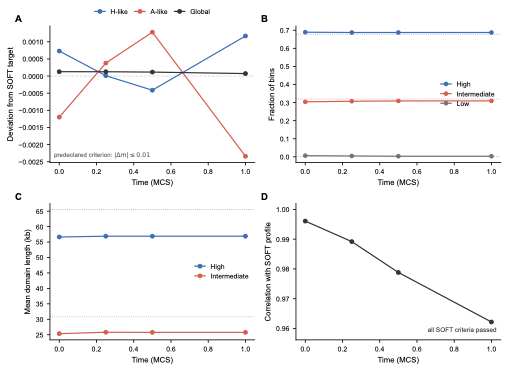

**Supplementary Figure S10. Finite-time preservation of the SOFT methylation architecture.** The frozen primary model was independently evaluated in the SOFT condition using 500 stochastic realizations and a fresh master seed. (A) Deviation of H-like, A-like, and global mean methylation from their SOFT reference values over 0–1 MCS. (B) Fraction of bins classified as high (m ≥ 0.60), intermediate (0.20 < m < 0.60), or low (m ≤ 0.20) methylation. Dotted horizontal lines denote the corresponding SOFT reference fractions. (C) Mean run length of high- and intermediate-methylation domains; dotted lines denote the empirical SOFT targets used to construct the reference architecture. (D) Pearson correlation between the simulated ensemble methylation profile and the initial SOFT profile. The predeclared preservation criteria required absolute errors ≤0.01 for H-like, A-like, and global means, ≤0.05 for high- and intermediate-state fractions, ≤25% relative error for mean high- and intermediate-state run lengths, and ≤10 kb absolute error for the corresponding medians. All criteria passed through the 1-MCS readout.

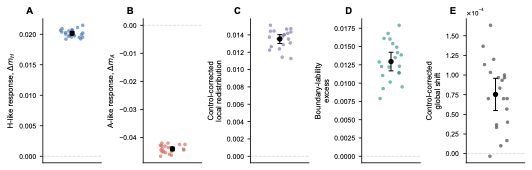

**Supplementary Figure S11. Reproducibility of the confinement response across independent master seeds.** The frozen model was repeated with 20 independent master seeds, each using 250 stochastic realizations per SOFT, independent SOFT control, and STIFF branch. (A) Control-corrected H-like methylation response, Δm_H. (B) Control-corrected A-like methylation response, Δm_A. (C) Control-corrected local redistribution, defined as the mean absolute STIFF–SOFT methylation change minus the corresponding value in the independent SOFT control. (D) Boundary-lability excess, defined as the control-corrected difference in absolute methylation change between bins ≤50 kb from a strong SOFT methylation-state boundary and bins ≥250 kb from a boundary. (E) Control-corrected global mean methylation shift. Colored points denote independent master seeds; black points and error bars show the mean and 95% confidence interval across master seeds. Dashed horizontal lines indicate zero response. The expected H-like gain, A-like loss, positive local redistribution, and positive boundary-lability excess occurred in all 20 master seeds, while the global mean shift remained small.

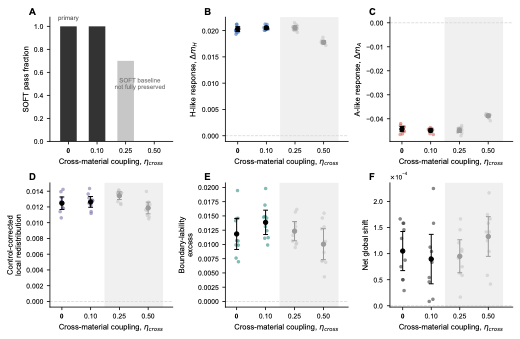

**Supplementary Figure S12. Sensitivity to cross-material cooperative coupling.** The fraction of the cooperative interaction J applied between different model material classes was varied as η_cross = 0, 0.10, 0.25, and 0.50 without recalibrating any other parameter. The primary model uses η_cross = 0. Each value was evaluated using 10 independent master seeds and 200 stochastic realizations per branch. (A) Fraction of master seeds satisfying all predeclared SOFT baseline-preservation criteria. (B) Control-corrected H-like response. (C) Control-corrected A-like response. (D) Control-corrected local redistribution. (E) Boundary-lability excess. (F) Raw net global mean methylation shift between STIFF and SOFT. Colored points denote individual master-seed estimates for parameter values at which the SOFT baseline remained fully valid; gray points and shading denote values for which complete SOFT preservation was not achieved and are shown as diagnostic sensitivity tests. Summary symbols show the mean and 95% confidence interval across master seeds. SOFT preservation was complete at η_cross = 0 and 0.10, decreased to 7/10 seeds at η_cross = 0.25, and failed in all seeds at η_cross = 0.50. These results indicate that the mechanism is robust to weak cross-material coupling but requires strongly preferential same-material cooperativity.

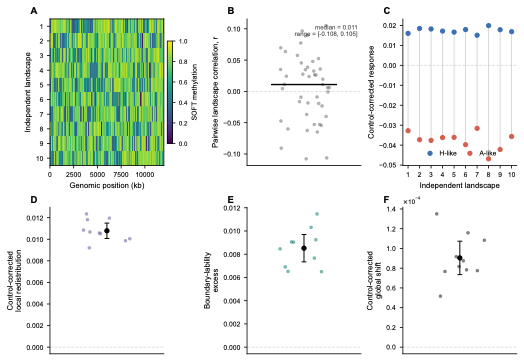

**Supplementary Figure S13. Generalization across independent empirical-like SOFT methylation landscapes.** Ten independent 12-Mb SOFT methylation landscapes were generated at 10-kb resolution using the same experimental methylation-state fractions and run-length constraints but different spatial arrangements. No landscape-specific model parameters were fitted. (A) Ensemble methylation profiles of the 10 starting landscapes. (B) Pairwise Pearson correlations among the 10 starting landscapes (45 landscape pairs), showing spatial independence; the median correlation was 0.011 with a range from −0.108 to +0.105. (C) Paired control-corrected H-like and A-like responses for each landscape; gray lines connect responses from the same landscape. (D) Control-corrected local redistribution. (E) Boundary-lability excess. (F) Control-corrected global mean methylation shift. Each landscape was evaluated using five independent master seeds and 200 stochastic realizations per branch. All 50 landscape-by-seed SOFT baseline validations passed. For panels C–F, effects were first averaged across the five master seeds within each landscape; the 10 landscape means therefore constitute the independent replicate unit. Points denote independent landscapes; black summary symbols and error bars denote the across-landscape mean and 95% confidence interval.

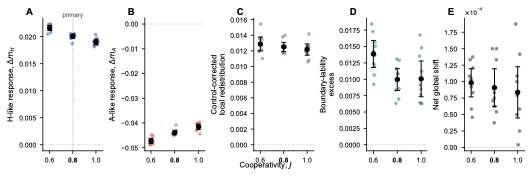

**Supplementary Figure S14. Robustness to cooperative coupling strength J.** The confinement response was evaluated at J = 0.6, 0.8, and 1.0 without recalibrating the local fields, μ, η_cross, γ, contact ranges, or readout time. The primary model uses J = 0.8. (A) Control-corrected H-like response. (B) Control-corrected A-like response. (C) Control-corrected local redistribution. (D) Boundary-lability excess. (E) Raw net global mean methylation shift between STIFF and SOFT. Each J value was evaluated using 10 independent master seeds and 200 stochastic realizations per branch. Colored points denote individual master-seed estimates; black points and error bars show the mean and 95% confidence interval across master seeds. Dashed horizontal lines indicate zero response, and the primary value is marked in the figure. All three J values satisfied the SOFT baseline-preservation criteria and retained the same directional confinement response.

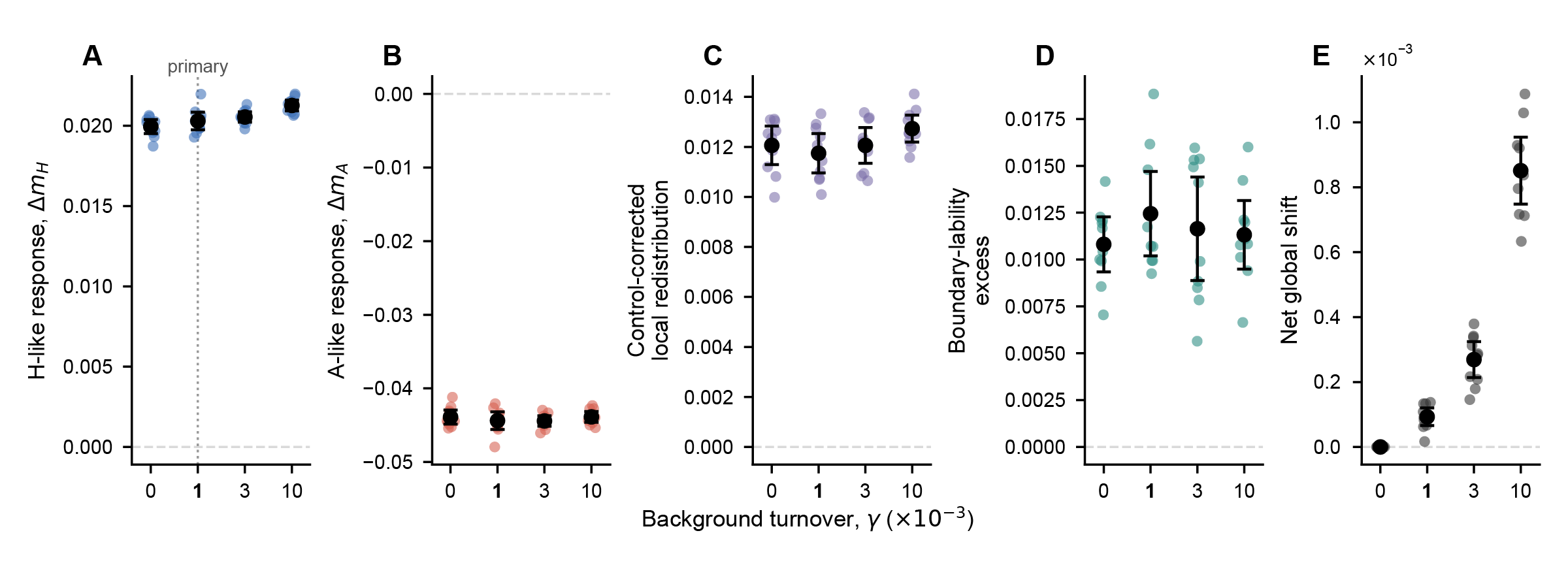

**Supplementary Figure S15. Robustness to background turnover rate γ.**The background turnover rate was varied as γ = 0, 0.001, 0.003, and 0.01 bin⁻¹ MCS⁻¹ without recalibrating any other model parameter. The primary model uses γ = 0.001. (A) Control-corrected H-like response. (B) Control-corrected A-like response. (C) Control-corrected local redistribution. (D) Boundary-lability excess. (E) Raw net global mean methylation shift between STIFF and SOFT. The four discrete γ values are plotted with equal spacing and labeled in units of ×10⁻³. Each value was evaluated using 10 independent master seeds and 200 stochastic realizations per branch. Colored points denote individual master-seed estimates; black points and error bars show the mean and 95% confidence interval across master seeds. Dashed horizontal lines indicate zero response, and the primary value is marked in the figure. All tested values satisfied the SOFT baseline-preservation criteria. The H-like gain, A-like loss, local redistribution, and boundary-lability response remain present at γ = 0, showing that the confinement response does not require the generic background turnover process.

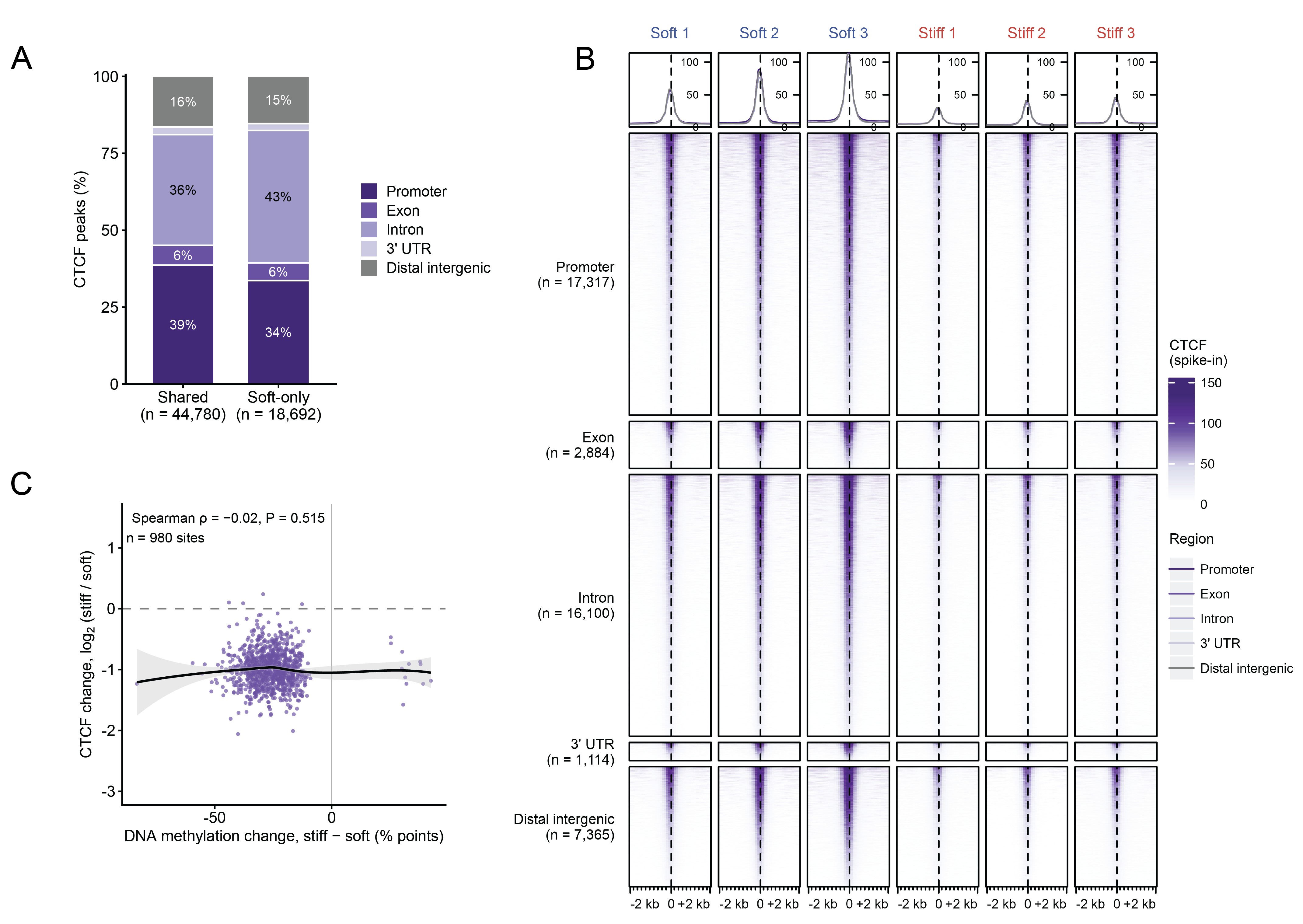

**Supplementary Figure S16**

**Supplementary Table T1.** Formulations of artificial ECM hydrogels.

| **Hydrogel** | **Collagen (mg/mL)** | **VLVG**  **alginate**  **(%**  **w/v)** | **CaCO_3_**  **(%**  **w/v)** | **GDL (mM)** |
| --- | --- | --- | --- | --- |
| **Soft viscous** | 4 | 1 | 0.10 | 40 |
| **Stiff viscous** | 4 | 1 | 0.30 | 120 |

**Supplementary Table T2.** Pharmacological inhibitors

| **Name** | **Synonyms** | **Target** | **Concentration** | **Manufacturer** |
| --- | --- | --- | --- | --- |
| Pam3CSK4 | TLR2 ligand | TLR2 | 1 μM | Invivogen |
| Poly(I:C)HMW | TLR3 ligand | TLR3 | 10 ug/mL | Invivogen |
| LPS | TLR4 ligand | TLR4 | 1 ug/mL | Invivogen |
| Decitabine | DNMT  inhibitor | DNMT | 0.5, 1 μM | Selleckchem |
| Blebbistatin |  | NMII | 25 μM | Selleckchem |
| ROCK  inhibitor | Y-27632 | ROCK | 25 μM | Selleckchem |

**Supplementary Table T3.** Details of primary, secondary antibodies and stains used in the study

| **Antibody** | **Specification** | **Dilution** | **Source** |
| --- | --- | --- | --- |
| Phalloidin i-Fluor |  | 1:1000 | Abcam (ab176753) |
| Lamin B1 | Rabbit Polyclonal | 1:400 | Proteintech (12987-1) |
| Phospho RelB (Ser552) | Rabbit Polyclonal | 1:200 | Cell Signaling Technologies (4999S) |
| H3K9me3 | Rabbit monoclonal | 1:400 | Invitrogen (720093) |
| Acetyl-Histone H3 (Lys27) | Rabbit monoclonal | 1:200 | Cell Signaling Technologies (8173) |
| DNMT3A | Rabbit monoclonal | 1:800 | Cell Signaling Technologies (32578) |
| CTCF | Rabbit monoclonal | 1:50 | Cell Signaling Technologies (3418) |
| DAPI |  | 1:2500 | Abcam (ab228549) |
| Secondary antibody | Alexa Fluor-594 (Goat anti-rabbit) | 1:300 | Molecular Probes (#A11037) |

### **Supplementary Table T4**. Parameters of the computational methylation model

Primary parameter values used in the final model.

| **Quantity** | **Final model value** |
| --- | --- |
| Genomic lattice | N = 1,200 bins; 10 kb per bin (12 Mb total) |
| Cooperative coupling | J = 0.8 |
| Cross-material coupling | 0 |
| Local fields | +0.23428517 (H-like); −0.23428517 (A-like); μ = −0.19635939 |
| Contact kernel | SOFT λ = 2; STIFF λ = 5 bins; cutoff = 4λ |
| Inverse-temperature scale | β = 1.0 |
| Background turnover | γ = 0.001 bin⁻¹ MCS⁻¹ |
| Simulation exposure and readout | N exchange proposals per MCS; t = 1 MCS |

**AUTHENTICATION OF KEY BIOLOGICAL AND/OR CHEMICAL RESOURCES**

1) All acquired compounds and reagents were authenticated for both identity and purity, based on certificate of analysis from manufacturers.
2) The fluorescent probes, laser and filter parameters related to the confocal microscopy have been extensively tested and selected to minimize potential cross talk between different excitation and emission channels
3) Primary human cells for in vitro experimentation were routinely tested for mycoplasma and bacterial contaminations.
